# FAIMS-GPF XL-MS: crosslinking-mass spectrometry based on gas-phase fractionation

**DOI:** 10.1101/2025.05.23.655748

**Authors:** Hugo Gizardin-Fredon, Salvatore Terrosu, Simon Pichard, Dursun Korkut, Cathy Braun, Arnaud Poterszman, Clément Charenton, Sarah Cianférani

## Abstract

Protein-protein interactions (PPIs) underpin nearly all cellular processes; therefore, mapping PPI and protein-protein association networks is critical for understanding how biological systems function in health and disease. However, many functionally relevant PPIs are transient and mediated by weak affinity interactions, making them challenging to detect. Using chemical cross-linking together with mass spectrometry (XL-MS) has emerged as an important category of methods for mapping PPIs, including those that are more dynamic and transient, although XL-MS workflows face limitations which hinder their routine use, especially in proteomic platforms. Here, to address these limitations, we developed a XL-MS workflow based on gas-phase fractionation (GPF) using high-field asymmetric waveform ion mobility spectrometry (FAIMS). Our optimized FAIMS-GPF XL-MS protocols exhibited improved performance when handling both low-complexity samples, such as purified Cdk7-Activating Kinase (CAK) heterotrimer, and high-complexity cross-linked HeLa lysates. In HeLa lysate, we identified 1,278 cross-links accounting for 213 PPIs without any in-solution fractionation, starting from less than 20 µg of sample. The method also proved effective for analyzing low-abundance, high-complexity samples such as spliceosomes. Overall, FAIMS-GPF enhanced the detection of cross-linked peptides and improved PPI coverage, thus offering a scalable, high-sensitivity solution for XL-MS integration with structural biology and functional proteomics applications.

## MAIN

Protein-protein interactions (PPIs) are essential drivers of cellular processes and function, including signal transduction, gene expression, cellular metabolism, cell growth, proliferation, and apoptosis. In general, proteins rarely function in isolation, their ability to interact with other proteins to form specific complexes and PPI networks is essential for ensuring coordinated execution of cellular tasks^1–4^. Therefore, identifying and characterizing PPIs is of critical importance for understanding fundamental cellular processes and biological pathways, as well as how their dysregulation contributes to disease mechanisms and pathogenesis. Although the exact number of PPIs formed in any given human cell remains unknown, several recent efforts have yielded information on tens of thousands of PPIs^5^. For example, Human Reference Interactome (HuRI) includes approximately 53,000 high-quality binary PPIs^6^, and hu.MAP 2.0 contains information on more than 57,000 predicted PPIs encompassing close to 10,000 human proteins^7^. However, based on some estimates, even these resources cover only a small fraction of all the PPIs that govern cellular function^8^. Therefore, deciphering PPIs and their networks remains of high interest and importance.

Currently, the most powerful methods for mapping PPIs and human interactome (*i.e.* the collection of all PPIs in a given cell) are based on the use of mass spectrometry (MS). In this context, the application of affinity-purification (AP) MS^9^, whereby high-affinity baits are used to capture PPIs, has been especially powerful in identifying interactions between human proteins, most recently resulting in capturing more than 100,000 interactions among approximately 14,500 proteins^10^. However, AP-MS based strategies underperform when detecting transient and low-affinity interactions, which are key determinants of cell signaling. On the other hand, although methods such as enzyme-catalyzed proximity labeling strategies (e.g. BioID, TurboID, APEX)^11–14^ coupled with MS are able to capture more transient interactions, they also have notable limitations including the need to fuse proteins of interest with enzymes that generate reactive species needed to label the interaction partners. Additionally, the labeling radius in these experiments varies between 10 – 20 nm^9,15^, which may lead to off-target reactivity and false positives.

Cross-linking mass spectrometry (XL-MS) represents a distinct set of strategies for identifying PPIs and defining their interfaces that uses chemical cross-linkers to form covalent linkages between interacting proteins before subjecting them to MS analysis^16^. The method is versatile and can be used with purified protein complexes^17^, organelles^18^, cells^4^, and even tissues^19^. XL-MS provides several advantages over the methods mentioned above, such as the ability to directly capture PPIs in their native environment without a priori, thus preserving physiologically relevant interactions. Additionally, XL-MS does not require the use of affinity reagents or fusion proteins and tags, and the availability of different XL cross-linkers with wide range of chemical reactivity profiles make these methods versatile and compatible with wide range of samples^16^. However, the XL reactions give low final yields, which leads to low intensity for XL-peptides in MS scans and limits PPI detection to high abundance proteins and stable interactions^20,21^. These effects amplify in case of higher complexity samples such as organelles or whole cells lysates.

In general, the performance of XL-MS for PPI mapping and interface definition in complex samples can be improved by using different strategies to reduce sample complexity as the use ofMS-cleavable XL reagents, such as sulfoxide-based DSBU^22^ and DSSO^23^ or enrichable reagents such as Protein Interaction Reporters (PIR)^24^, Azide-A-DSBSO^25^ or phosphonate-based PhoX^26^. The most extensive studies, although essential for uncovering a high number of PPIs, often rely on the use of off-line chromatographic fractionation^22,27–29^ with the collection of numerous fractions (>20), which requires substantial amounts of starting material (>500 µg) and significantly increases instrument time. During past years, ion mobility techniques such as high-field asymmetric-waveform ion-mobility spectrometry (FAIMS) and trapped ion mobility spectrometry (TIMS) also emerged as a key strategy to improve peptides identifications across various fields of proteomics. These techniques offer an additional separation dimension in gas-phase separation that is strongly influenced by peptides charges. In this later context, given that cross-linked peptides typically carry higher charge (≥ 3+) compared to linear peptides (+2), the use of FAIMS (High-field asymmetric-waveform ion-mobility spectrometry)^30,31^ in XL-MS workflows either with MS-cleavable reagents^30–32^ or enrichable PhoX^33^ resulted in significant improvement of XL-IDs when combined with in-solution fractionation strategies. However, the use of FAIMS as a standalone gas-phase fractionation (GFP) method to fully replace in-solution fractionation has not yet been evaluated and may further enhance our ability to harness the power of XL-MS for identifying protein–protein interactions (PPIs) in complex systems..

Here, we developed a workflow that employs FAIMS-GPF, coupled to a fast and sensitive sample preparation using DSSO^23^ and automated peptide cleanup for improved XL-MS based identification of PPIs. As a proof-of-concept, we applied this method for several case studies of different systems, a structural biology project focusing on Cdk7-Activating Kinase (CAK) trimer, a proteome wide study of cross-linked HeLa lysate, and a study of PPIs in a challenging spliceosome sample where FAIMS-GPF was combined with mass photometry (MP). We demonstrated that FAIMS-GPF XL-MS approach we described achieves unprecedented sensitivity and throughput for all sample complexities. Furthermore, we showed that eliminating in-solution fractionation enables the use of a consistent, adaptable and streamlined workflow across all sample types.

## RESULTS

### General description of the FAIMS-GPF XL-MS workflow

To streamline and increase efficacy of the workflow, we optimized our sample preparation step to a 1-day process that uses MP for quality control step prior to a 45-minute cross-linking reaction with DSSO, overnight Trypsin/LysC digestion and an automated solid-phase extraction (SPE) sample clean-up (**Fig. 1a**). The samples were then subjected to nano-scale liquid chromatography (nLC) prior to FAIMS filtering and mass spectrometry (MS) (**Fig. 1b**). Briefly, FAIMS uses an asymmetric waveform that alternate between a high-field voltage (called dispersion voltage, DV) of one polarity and a low field voltage of the opposite polarity^34,35^. If the ion has a difference in mobility between high and low field (due to charge for example), it collides on one electrode and is not detected in MS. An additional direct current called “Compensation Voltage” (CV) allows compensating this ion displacement to bias the transmission of a subset of ions of interest ^34,35^, which aims in our case at enriching cross-linked peptides in gas-phase^35,36,31^.

**Figure 1.**
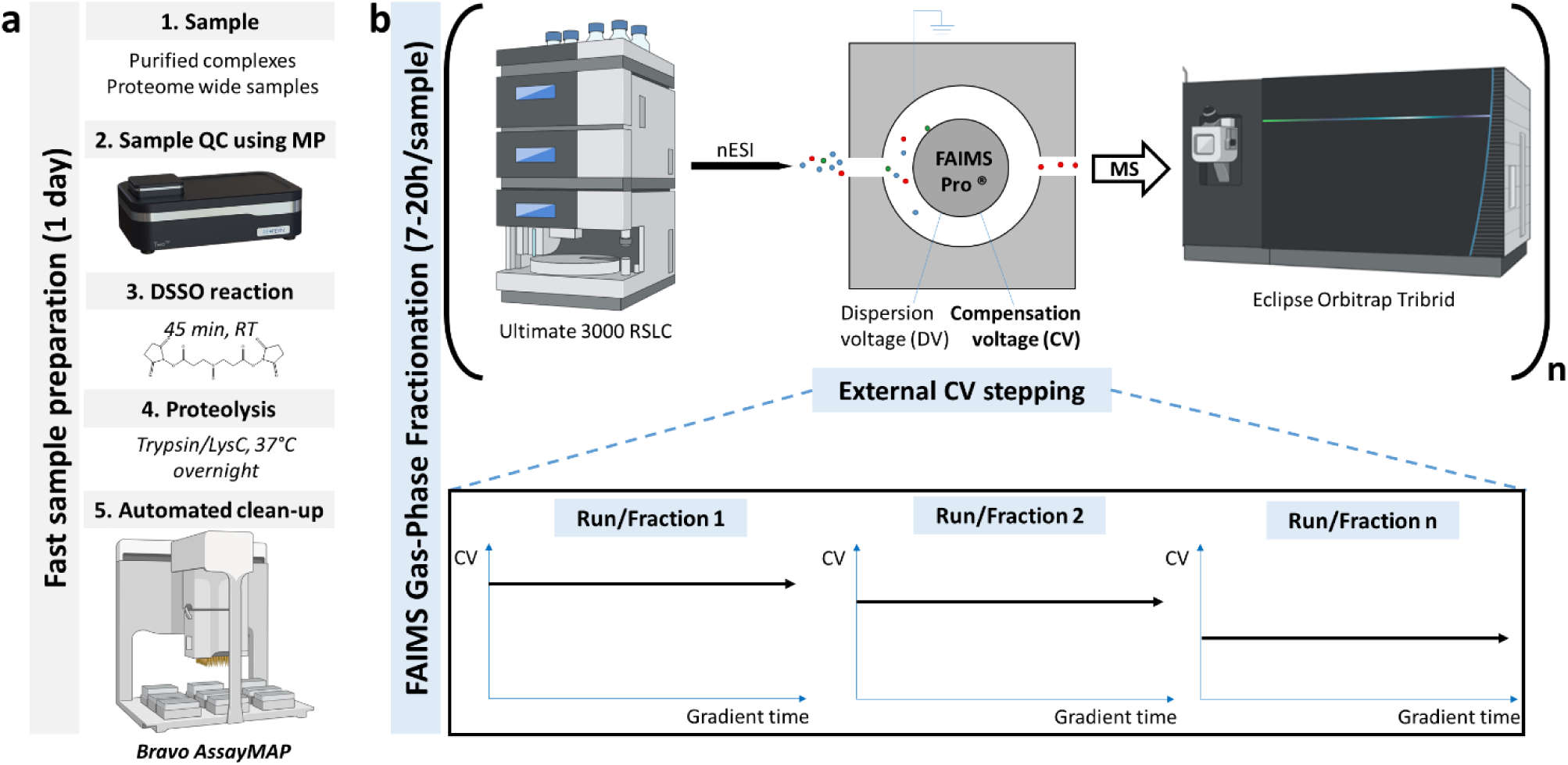
Experimental design for the evaluation of FAIMS-GPF in XL-MS workflows. **(a)** We applied a fast sample preparation step (1 day). After a mass photometry QC, samples are submitted to XL reactions using DSSO, overnight digested with Trypsin/LysC and cleaned up using an automated solid-phase extraction (SPE) method on the Bravo AssayMAP robot. **(b)** For FAIMS-GPF, 400 ng are injected in successive LC-FAIMS-MS/MS runs, each time with a different compensation voltage (CV) in a so-called ‘external stepping’ fashion. We started by screening different values before selected the best associations for our optimized methods on each sample.

As the first step in developing our workflow, we evaluated two different FAIMS-based GPF modes using either (i) “internal CV stepping”, where a single LC gradient is followed by different successive compensation voltage (CV) values that alternate with a defined cycle time; or (ii) “external CV stepping” where analysis is done using several successive LC-FAIMS-MS/MS runs each with a single CV value^37^ (**Supplementary Figure 1a**). We confirmed that external CV stepping with a single tailored CV provides higher numbers of XLs than internal stepping (**Supplementary Fig. 1b**), in agreement with published data^37,38^. Therefore, our optimized FAIMS-GPF XL-MS workflow includes a cross-linking step using MS-cleavable DSSO, a liquid digestion (Trypsin/LysC), and a fast automated peptide cleanup (*i.e.* fast sample preparation process; **Fig. 1a**), followed by a nLC-FAIMS-GPF-MS/MS with external CV stepping (**Fig. 1b**). This workflow was then evaluated and optimized on two samples of increasing complexity, the ∼110 kDa trimeric CAK complex and ‘native’ HeLa cell lysate, as described below.

### Using FAIMS-GPF XL-MS to extend structural biology insights into CAK complex

To evaluate performance of FAIMS-GPF when applied to the low complexity system, *i.e.* a purified protein complex, we used ∼110 kDa trimeric CAK (CdK7-Activating Kinase) complex, composed of CDK7, MAT1 and Cyclin H. CAK is involved in the regulation of cell cycle and transcription processes either alone or as a component of the general transcription factor II H (TFIIH)^39–41^. As the first step, we screened different CV conditions to identify voltage values that yield the largest number of unique XLs. We pooled three independent XL reactions each starting from 20 µg of CAK complex before digestion and peptide cleanup steps. We then directly analyzed the pooled sample using nanoLC-FAIMS-MS/MS (400 ng injected per CV run) with a gradient length of 105 min (90 min separation) to afford reasonable throughputs. The screening of CV values was done using external CV-stepping with steps of 5 V from -40 V to -90 V (11 CV values, **Fig. 2a**) and was compared to unfractionated XL-CAK. The FAIMS CV screening showed that single nanoLC-FAIMS-MS/MS runs with voltages ranging from -50 V to -70 V were all able to increase the total number of XL identifications. The highest number was obtained at -55 V (**Fig. 2a**), with 60 unique XLs identified (compared to 33 unique XLs for unfractionated CAK). This is most likely due to the “gas-phase enrichment” of ≥ 4+ charge states in the CV range -50 to -70 V, with CV of -55 V yielding up to 72 % of ≥ 4+ precursors (**Fig. 2b**). We evaluated the potential of cumulative external FAIMS-GPF steps using successive runs (‘n-CV GPF’), and observed that although increase in unique XL pairs identified was evident even with 1-CV GPF, 4-CV GPF method allowed 97 % coverage of all unique XL pairs, with that number changing insignificantly with further increase to 11-CV GPF (**Fig. 2c**). Thus, 4-CV GPF setup provides almost complete coverage of unique XL pairs while only requiring 7 h of instrument time and a total of 1.6 µg of starting material; it also almost triples the number of unique XLs identifications (x2.6) compared to analysis without FAIMS (**Fig. 2d**).

**Figure 2:**
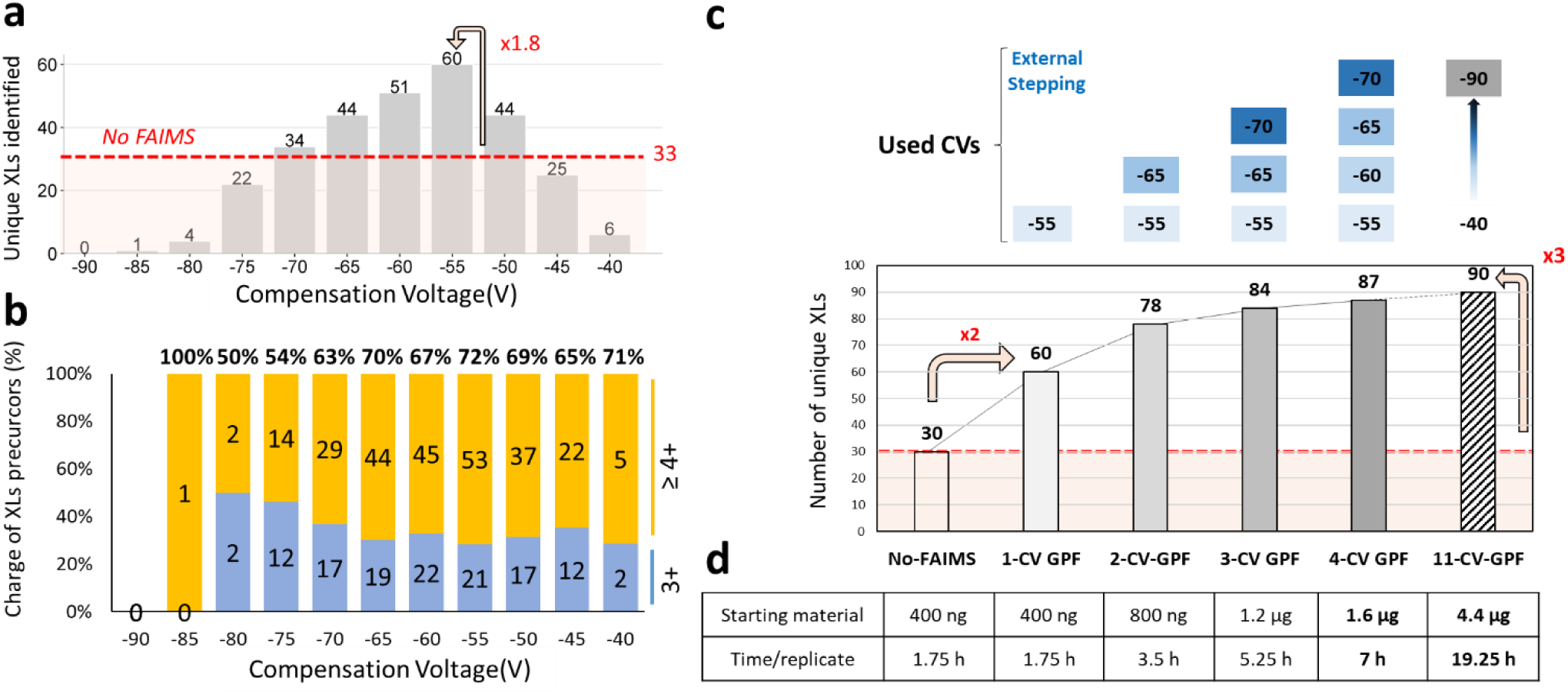
Screening of Compensation Voltage values on cross-linked CAK (5 V external stepping, n=1 injection per CV). **(a)** Number of identified unique cross-links per CV value. **(b)** Charge repartition of cross-link spectrum matches (CSM) precursors per CV value with 3+ in blue and ≥ 4 + in orange with the corresponding umber of precursors indicated inside bars. The percentage of ≥ 4+ charges is indicated on top of the columns. **(c)** Number of unique XL identified as a function of the number of CV-values chosen. **(d)** The table summarizes the total amount of CAK complex consumed and the associated time of LC-MS/MS analysis.

We applied 4-CV GPF protocol to a “real life” XL-MS experiment on CAK complex in three independent XL replicates. We first performed native mass photometry (nMP) and native mass spectrometry (nMS) to assess the quality of our sample and checked the integrity of the CAK heterotrimer, before XL-MS experiments. Native MS revealed presence of up to three phosphorylation on CAK trimer (**Fig. 3a**), with the proteoform bearing a single phosphorylation as a predominant form (MW_exp_ = 113,877 ± 3 Da, MW_theo_ = 113,879 Da). nMP allowed us to further confirm the main species (>95 % of sample) as intact CAK trimer (116.1 ± 1.0 kDa, **Fig. 3a**). To monitor the outcome of the DSSO XL reaction, we further used denaturing MP^42^ and confirmed that between 60 and 70 % of CAK trimer was covalently stabilized in each XL-replicates (**Fig. 3b, Supplementary Fig. 2**). Additional species were also identified corresponding to free monomers (16 ± 8 %, 10 ± 8 %, 7. ± 1 % for XL replicates 1, 2, 3), dimers (14 ± 3 %, 13 ± 11 %, 16 ± 2 % for XL replicates 1, 2, 3), as well as a low abundance of dimers of CAK trimer (5 ± 2 %, 11 ± 2 %, and 8 ± 1 % for XL replicates 1, 2, 3). We next analyzed the XL-replicates with the optimized 4-CV GPF method and compared it to analyses without FAIMS. Considering only unique XLs pairs present in at least 2 out of 3 replicates (**Supplementary Fig. 3**), a final validated dataset of 48 XLs was obtained without FAIMS (**Supplementary Table 1**), the 4-CV GPF protocol identified 82 XLs (**Supplementary Table 2**). When comparing the two datasets, 40 XL pairs (44 %) were found in common, while 43 XLs (47 %) were specific to the 4-CV GPF protocol and only 8 XLs (9 %) were uniquely found in the unfractionated run (**Fig. 3c-e**). Importantly, the 4-CV FAIMS-GPF protocol allowed us to identify additional interacting regions that have not been detected without FAIMS, such as interactions in the coiled coil domain of MAT1 (binding to ARCH) and the C-terminal of kinase domain of CDK7 (**Fig. 3d**; **Supplementary Fig. 4**).

**Figure 3:**
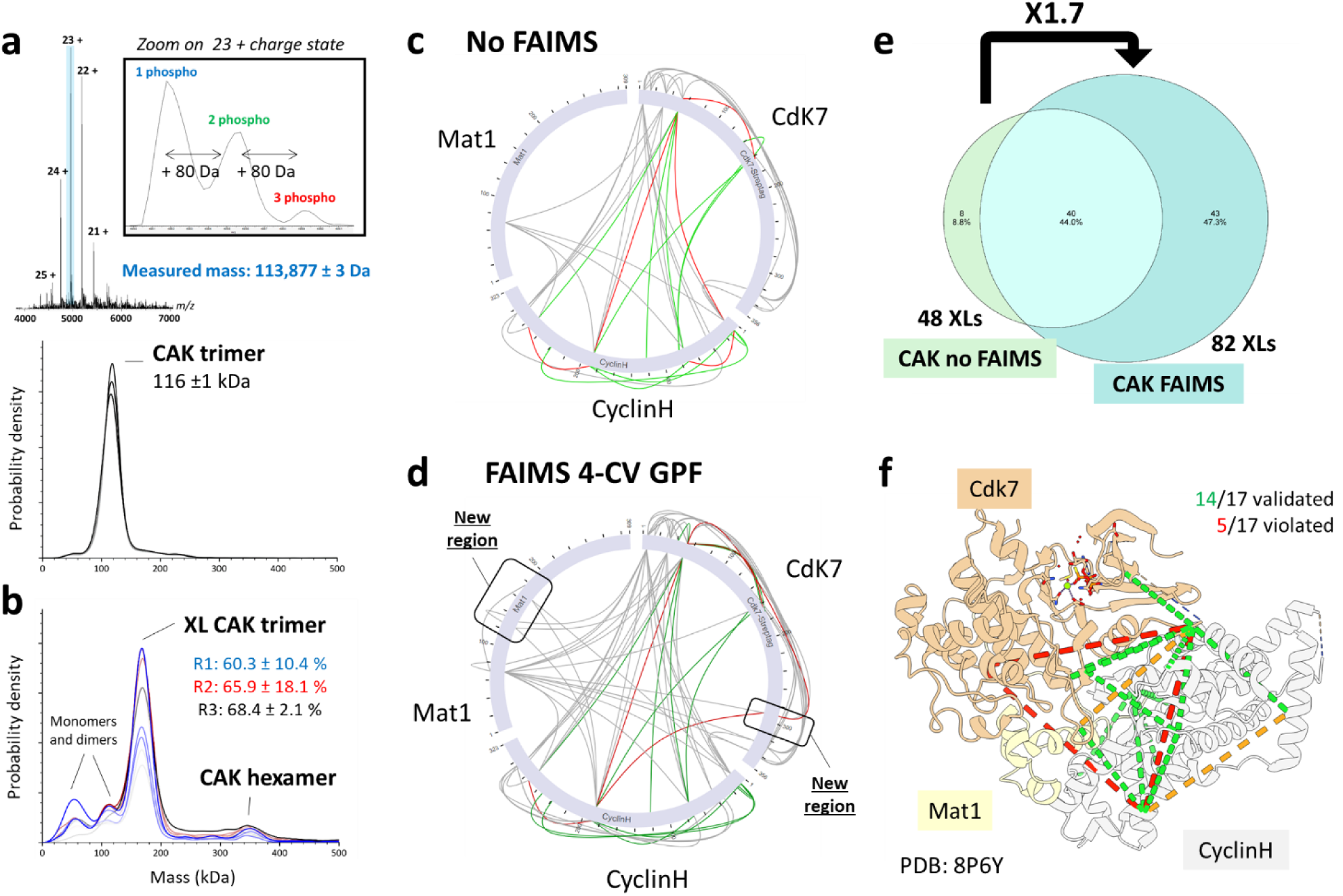
Comparison of XL-MS results in triplicates on CAK complex using classical method and FAIMS-4-CV GPF. **(a)** Quality control of CAK sample before cross-linking reaction using native-MS confirms the presence of CAK trimer with 1 (113,877 ± 3 Da), 2 or 3 phosphorylation. nMP allows to quantify > 95 % of intact CAK in the sample, with measured mass of 116 ± 1 kDa. **(b)** dMP control of cross-linked CAK sample triplicate using dMP shows the stabilization of trimer and the co-existence of free monomers and sub-complexes as well as a minor population of dimers of CAK heterotrimers. Measurement repeats of XL replicates 1, 2, 3 are color-coded respectively in shades of blue, shades of black and shades of red. Mass shift between theory and XL-CAK measured mass in dMP corresponds to binding of mono-links and cross-links. Standard deviations are calculated with measurement repeats (n=3). **(c)** Circular plot displays identified XLs on CAK partners’ sequences for CAK XL-MS experiments without FAIMS and **(d)** CAK XL-MS experiments with 4-CV FAIMS GPF. Additional regions of interaction between are covered thanks to FAIMS-GPF in Mat1 and CdK7. **(e)** Venn diagram shows the overlap of validated unique XLs (file threshold 2/3) in experiments without and with FAIMS, which allows increasing by 70% the number of XLs identified and identify 43 new XLs. **(f)** We could map 19 XLs on Cryo-EM structure of CAK in complex with ATPgS (PDB: 8P6Y), out of which 14 were distance validated (74 %) with a strict Cα-Cα threshold of 30 Å (highlighted in green), five were distance-violated (orange and red).

We also examined how well the XLs we identified using the 4-CV FAIMS-GPF protocol agree with the recently published cryo-EM structure of CAK (PDB: 8P6Y, **Figure 3f**). Given that the density in the deposited CAK structure was incomplete, we could only map 19 (out of 82) XLs; nonetheless, majority of the XLs we could map were distance validated with a strict Cα-Cα threshold of 30 Å (14 out of 19; 74%), while only five were distance-violated. Looking beyond the cryo-EM structure, In particular, we identified 24 interactions between Cdk7 and Cyclin H, which involve residues scattered over the entire cyclin H sequence and mostly in the N-terminal region of Cdk7. We also observed 11 XLs between pairs of residues that could be modelled from X-ray structure of Cyclin H and Cdk7^43,44^. Interestingly, the XLs between Cdk7 and Cyclin H involve residues scattered over the entire Cyclin H sequence and mostly in the N-terminal region of Cdk7. These XLs are compatible with an AlphaFold-derived model^45,46^ obtained by docking the AlphaFold models of Cdk7 and Cyclin H onto the cryo-EM structure of CAK (**Supplementary Fig. 5**). The XL dataset also contains 17 XL-pairs composed of one peptide from either Cdk7 or Cyclin H and one from MAT1. All detected XLs involve a residue from MAT1 located in the N-terminal part of the protein, mainly in the helical bundle, which shows that this region is able to fold back on the Cdk7/Cyclin H dimer (**Supplementary Fig. 6**). Therefore, although the cryo-EM structures of CAK^47,48^ does not include density for the N-terminal part of MAT1 due to the intrinsic flexibility of this region with respect to the rest of the CAK complex, our XL data was able to capture the transients interactions this domain makes with Cdk7/Cyclin H dimer.

Taken together, our 4-CV FAIMS-GPF workflow allowed us to map additional interactions and interfaces involved in CAK complex. Importantly, the complete XL-MS experiment (done in triplicate) was carried in only 3.5 days (21 h of instrument time) with only 4.8 µg of sample. Thus, FAIMS-GPF XL-MS provides a significant increase in sensitivity without requiring increase in the amount of starting material consumed or instrument time.

#### Proteome-wide FAIMS-GPF XL-MS experiments in whole cell lysates

We next evaluated the potential of using FAIMS-GPF XL-MS on a higher complexity sample, *i.e.* a whole-cell native HeLa lysate. As in the previous case study, we first optimized FAIMS-GPF conditions by testing a wide range of CV voltage values, displaying a similar charge-related filtering, and evaluating the optimal number of successive FAIMS-GPF runs (**Fig. 4a-c, Supplementary figures 7**). Conversely to the low complexity CAK sample, the number of unique XLs identified does not plateau after 4-CV GPF and the increase is almost proportional to the number of gas-phase fractions used (R^2^ = 0.97, **Supplementary figures 8 and 9**). Ultimately, we determined that a protocol with 11-CV FAIMS-GPF allowed identification of 1250 unique XLs, which is close to five time the number of unique XLs identified without FAIMS (**Fig. 4d**). This significant increase in XLs identifications was achieved using less than 1 day of instrument time and consuming as low as 4.4 µg of sample per sample to analyze (*e.g.* per XL replicates).

**Figure 4:**
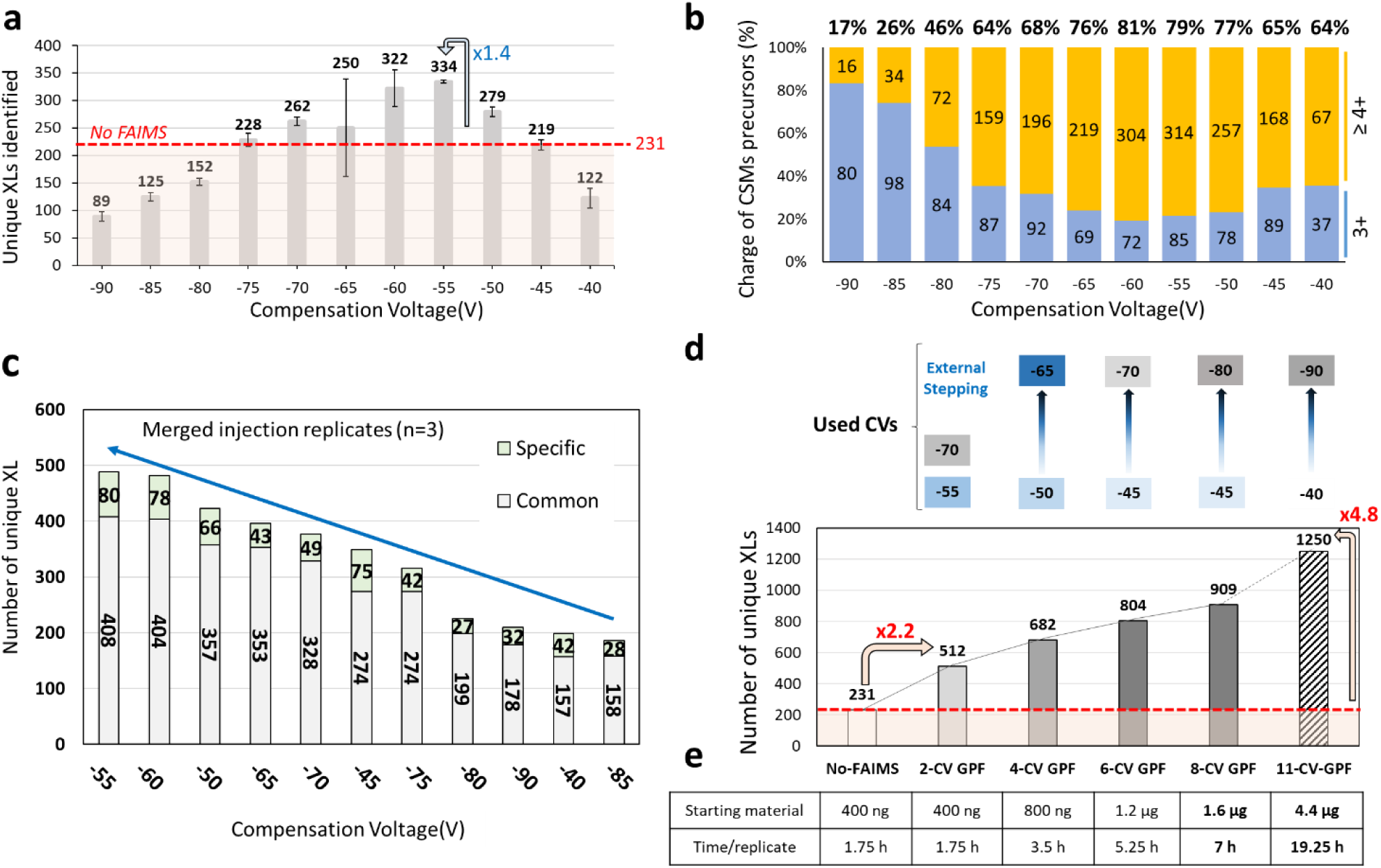
Evaluation of FAIMS-GPF approach on cross-linked HeLa lysates. **(a)** Screening of CV values showing the number of identified unique cross-links per CV value. Average ± SD correspond to injection replicates (n=3). **(b)** Average charge repartition of CSM precursors per CV value (n=3 injection per CV) with corresponding number of precursors indicated inside bars. Blue bars represent respectively 3+ and orange bars ≥ 4+ charge states. Percentage of ≥ 4 + charges states are written on top of bars. **(c)** According to the number of fractions needed (‘n-CV GPF’), CV values with the highest absolute number of unique XLs identified were successively associated, following order indicated with blue arrow (from merged injection replicates, n=3). This order is closely related to the number of XLs specific to the CV value. **(e)** Evaluation of the number of gas-phase fractions needed for XL-HeLa lysate and the potential throughput and sensitivity reached, using the first injection replicates.

To highlight the benefits of our FAIMS-GPF approach, we carried a complete XL-MS experiment by cross-linking three HeLa lysates replicates with 2 mM DSSO as a cross-linking reagent (**Fig. 5**). We compared results of 11-CV FAIMS-GPF protocol with results obtained using unfractionated HeLa lysate. While our benchmark experiment performed without FAIMS identified 443 unique XLs pairs (381 intra-molecular, and 62 inter-molecular XLs representing 59 PPIs), the 11-CV FAIMS-GPF method allowed us to identify 1269 XL pairs (1015 intra-molecular and 254 inter-molecular XLs, representing 213 PPIs) (**Fig. 5a**). This represents a 2.8 time increase in number of unique XLs identifications, with 915 new XLs, and a 3.6 time increase in PPIs with 178 new PPIs identified compared to FAIMS-free analysis (**Fig. 5a**, **Supplementary Fig. 10**). Importantly, the reproducibility of the FAIMS-GPF was also higher than without FAIMS (65.9 % vs. 43.3% without FAIMS, **Fig. 5b**, **Supplementary Fig. 11**).

**Figure 5:**
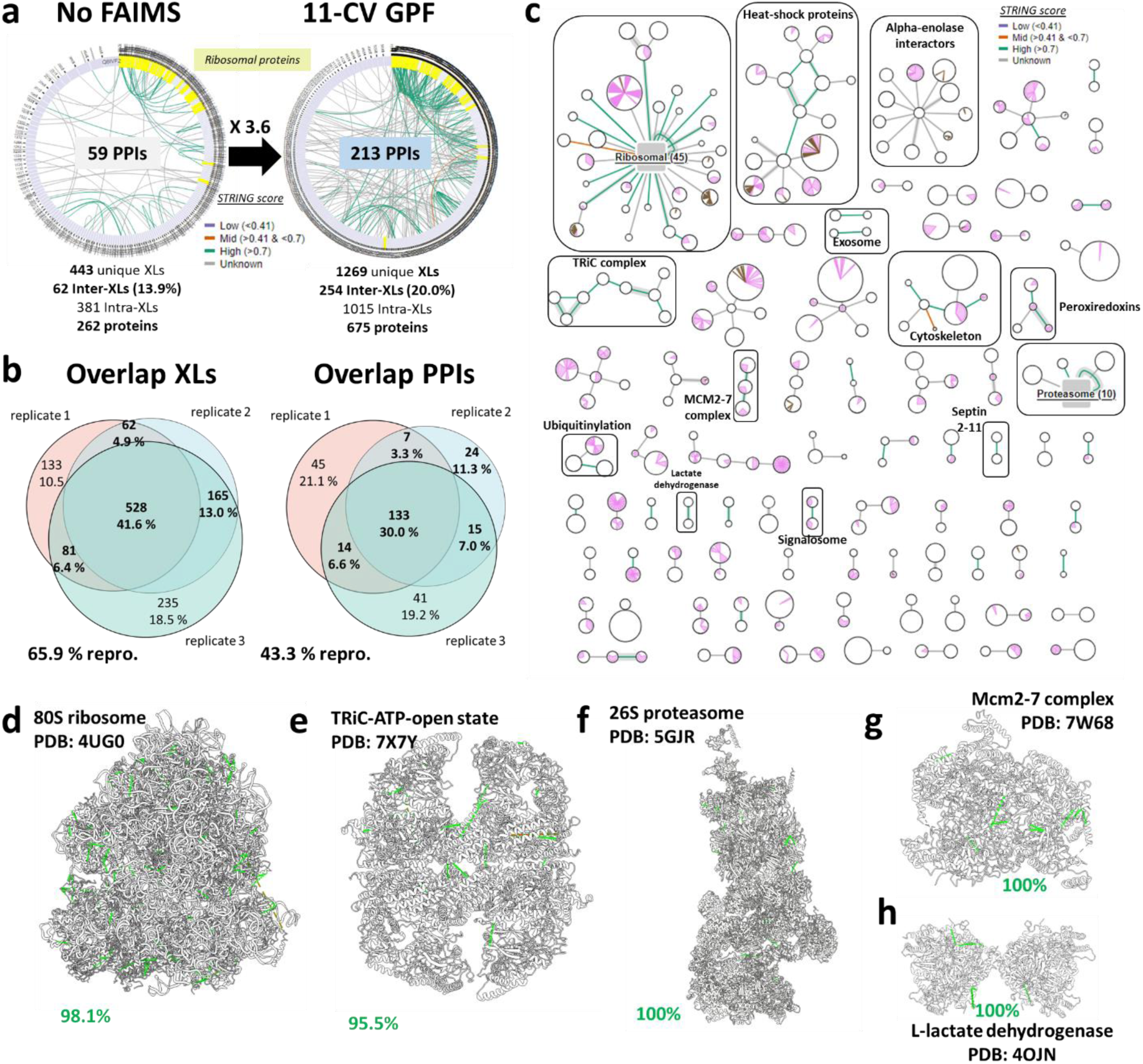
Results of XL-MS experiments using 11-CV FAIMS-GPF method on native HeLa Lysate cross-linked with DSSO in triplicate. **(a)** Circular plot compare the inter-protein XLs identified from three independent XL replicates without FAIMS or with the 11-CV GPF method. A 3.6 times increase in XLs identification is observed with FAIMS GPF leading to the identification of 1269 XLs in total accounting for 213 PPIs. Ribosomal proteins, highlighted in yellow are known to be a hotpot for interaction, and known interactions from STRING database are highlighted in green if highly confident (>0.7) or orange if of lower confidence (< 0.7). **(b)** Reproducibility (in ≥ 2/3 replicates) of unique XLs and PPIs identified in replicates (n=3) is high with respectively 65.9 % and 43.3 %. **(c)** PPIs network obtained after 11-CV GPF displays 213 PPIs, where ribosome and proteasome are hotspots for interactions (both collapsed). PPIs from major complexes or important biological functions are circled in black. Size of nodes is proportional to protein length with known domains (from Uniprot) highlighted in pink as well as transmembrane domains highlighted in brown. Thickness of edges is proportional to the XL number for each PPI and colour-coded according to their STRING score (green > 0.7, orange < 0.7). Identified intra-protein and inter-protein XLs could be mapped on known 3D structures of different complexes with different sizes: **(d)** 80S Ribosome**, (e)** TRiC-ATP-open state, **(f)** 26S proteasome, **(g)** Mcm2-7 complex and **(h)** L-Lactate dehydrogenase. Overall, 97.5 % of XLs were distance-validated.

**Figure 6:**
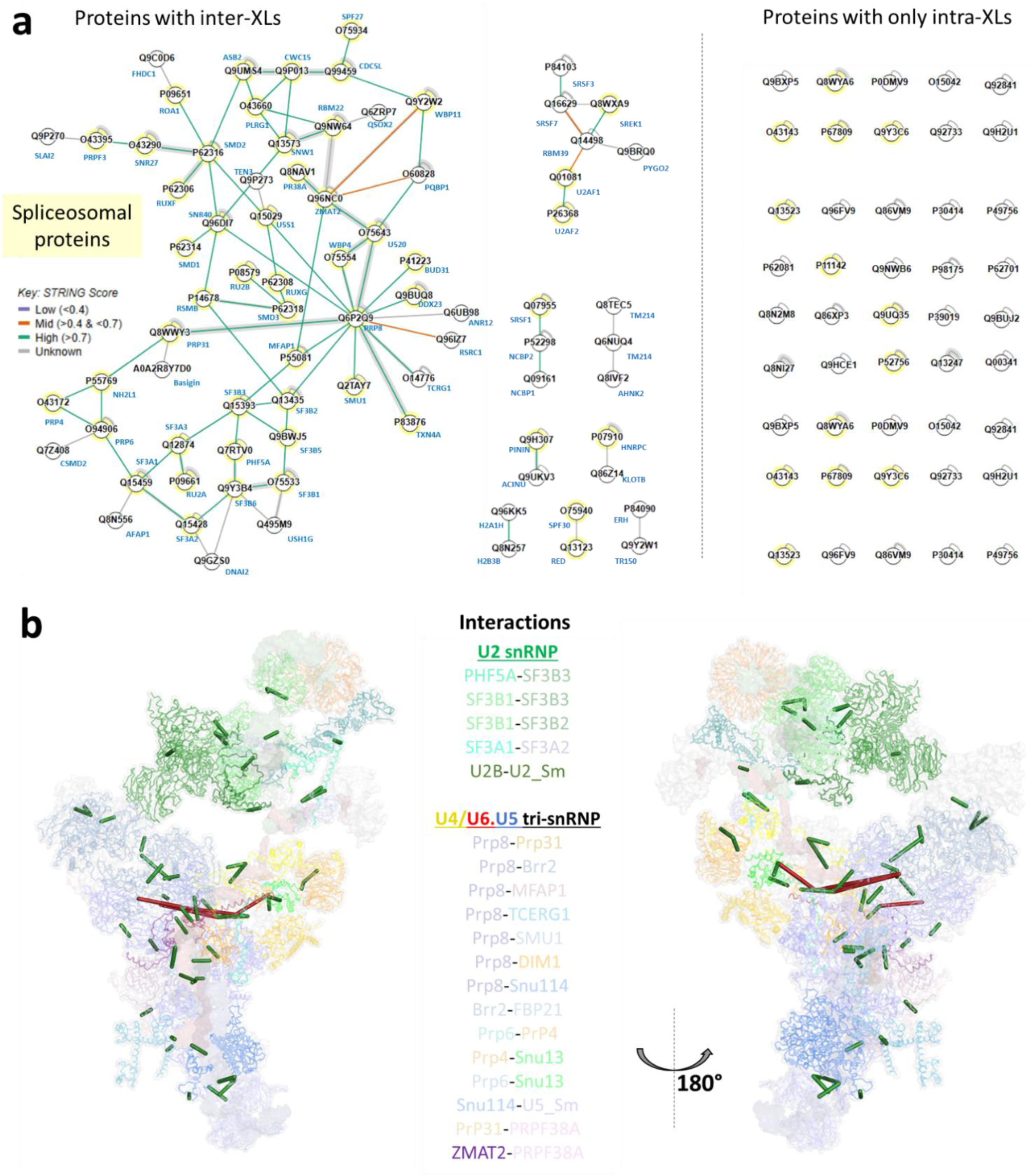
Results of XL-MS workflow with 11-CV FAIMS-GPF method to study spliceosomal proteins interactions. **(a)** xiView network shows the 283 reproducible XLs identified in at least 2/3 replicates after 11-CV-GPF experiments. Nodes represent cross-linked proteins among which the ones from spliceosome (Gene Ontology) are highlighted in yellow. Lines thickness is proportional to the number of XLs and interactions found in STRING database are color-coded in green for scores > 0.7 (confident) and in orange for scores <0.7. **(b)** Two different views of the human B-complex (PDB: 8QO9) show the spatial distribution of cross-links across the assembly. Intermolecular cross-links are mapped on the structure and highlighted as green (distance validated) or red (distance violated) lines according to distance between cross-linked residues (DSSO linker < 30 A). Proteins involved in intermolecular cross-linking are shown in ribbon and colored according to the color scheme shown in the legend, snRNA are shown as a colored surface: U2 in green, U4 in yellow, U5 in blue and U6 in red respectively.

To functionally annotate the unique XLs we identified, we performed a Gene Ontology (GO) enrichment analysis of the cross-linked proteins uniquely identified in FAIMS-GPF experiments (**Supplementary Fig. 12**). The enriched GO terms (p < 10E-20) corresponded to ‘cellular response to a stimuli’, ‘VEGFA/VEGFR signaling’, ‘60S ribosomal subunits’, ‘protein-RNA complex organization’, ‘regulation of transcription’, ‘neutrophil degranulation’, ‘carbon metabolism’, and ‘protein folding’. The highest number of PPIs was identified for ribosomal proteins (**Fig. 5c**; **Supplementary Fig. 14a**), 31 between ribosomal proteins and 21 between ribosomal proteins and other interactors, among which 8 were known interactions reported in STRING database^49^ (*e.g.* RL23-EF2, RS12-SEBP1, RL31-BTF3) and 13 not previously known ones (*e.g.* RL4-FAM47E, RS7-PIGG, RS6-PEPD). We also observed a number of PPIs for proteasome subunits (**Fig. 5c**; **Supplementary Fig. 14b**). Additionally, we retrieved PPIs from other important complexes such TRiC (8 PPIs), MCM 2-7 complex (2), lactate dehydrogenase (1), signalosome (1), or Septin 2-11 (1), and interactions related to different biological processes or functions as ubiquitination (2), heat-shock proteins (15) or cytoskeleton (6) (**Fig. 5c**). Despite using a whole cell lysate, no interactions were found between “non-compatible” cell compartments. Among identified PPIs, 136 were present in the STRING database and 86.7 % of them were of very high confidence (scores > 0.9) (**Supplementary Fig. 15a**), highlighting the high degree of agreement between our data and previously reported PPIs. As a further quality check, we mapped MS-detected intra- and inter-XLs on available 3D structures for five selected complexes of different sizes (**Fig. 5d-h**). In total, 159 XLs were distance-validated at 97.5 % across all complexes (**Supplementary Fig. 15b**).

Importantly, a significant portion of the PPIs we identified (118 out of 254) are not currently in the STRING database suggesting that FAIMS-GPF XL-MS has the ability to identify novel direct interactions. For example, we identified a set of previously undocumented interactions involving alpha-enolase (ENO1) and major heat-shock proteins (**Fig. 5c, Supplementary Fig. 16**). ENO1, a glycolytic enzyme overexpressed in various cancers, is known to support tumor progression through metabolic reprogramming, proliferation, and immune evasion^50,51^. Our data revealed potentially novel ENO1 interactions with ZNF521^52,53^ (a transcriptional regulator linked to cell differentiation), CKAP2^54^ and Coronin-2B^55^ (involved in cytoskeleton and actin dynamics), SNUT1^56^ (an RNA splicing factor), and membrane-associated proteins such as MYO1A^57^ and APLD1^58^. This suggests that ENO1 functions as a multifunctional hub bridging metabolism, gene regulation, and signaling in oncogenic contexts. In parallel, we identified potentially new interactions between HSPA1B, HSPA8, HSP90AB1, and HSPD1 with proteins like U2SURP^59^ (splicing), DIAPH1^60^ (cytoskeleton), and DNAJC13^61^ (vesicle trafficking), highlighting an expanded chaperone network that integrates proteostasis with broader cellular processes. Together, these findings reveal new layers of functional connectivity for both ENO1 and heat shock proteins, providing a valuable resource for exploring their roles in cancer and cellular homeostasis. Although newly identified PPIs remain to be experimentally validated and functionally characterized in future studies, these results highlight the potential of FAIMS-GPF XL-MS as a tool for discovering new biologically relevant interactions. One possible reason why we were able to identify novel PPIs may be the increased sensitivity of FAIMS-GPF compared to FAIMS-free experiments, as highlighted by label-free quantification (LFQ) experiments (**Supplementary Fig. 13**).

Taken together, nanoLC-FAIMS-MS/MS workflow we developed was able to identify 1269 XLs and 213 PPIs overall (almost 3 times the number of XLs obtained without FAIMS) in less than 3 days of instrument time and using only 5.5 µg per XL replicate (for a total < 20 µg). In comparison, similar performance using existing methods would require extensive in-solution fractionations (> 20 fractions) using high amounts of starting material (> 100 µg per sample, usually around 500 µg)^27,28,62–64^. Therefore, the 11-CV FAIMS-GPF protocol achieved an unprecedented sensitivity and high reproducibility in shorter time and using small amounts of starting material (as low as few micrograms). This improved performance allowed us to identify novel PPIs between important cellular proteins, suggesting that our method can be used to discover new aspects of cellular function.

### The 11-CV FAIMS-GPF protocol enables the study of challenging spliceosome complexes

Finally, we used the 11-CV FAIMS-GPF XL-MS method combined with MP on a challenging biological ribonucleoprotein sample, the spliceosome. During pre-mRNA splicing, the spliceosome removes non-coding introns and joins coding exons together to produce mature messenger RNAs (mRNAs)^65–67^. This megadalton-protein-RNA machinery (> 200 partners) forms *de novo* on each intron and is subjected to extensive compositional and conformational transitions during the splicing process^35–41^. Although several studies have highlighted the value of XL-MS, especially when combined with cryo-EM, in providing critical insights into the assembly and function of the spliceosome^69–73^, purifying sufficient amounts of material for informative XL-MS studies remains a challenge. For example, our procedures routinely yield < 40 µg of pre-catalytic spliceosome, which is not enough to perform extensive fractionations^28^ described for such samples and even worse to replicate XL reaction asrecommended for XL-MS of purified complexes^74,75^. Thus, we examined whether 11-CV FAIMS-GPF XL-MS method overcomes these limitations.

We verified sample integrity using nMP^76,77^, confirming the presence of intact RNA-bound spliceosomes (1749 ± 43 kDa) and two sub-complexes (1024 ± 124 kDa and 487 ± 13 kDa, **Supplementary Fig. 17a and b**). We split the sample in three, and carried XL reaction in triplicate (11 µg each per reaction) using 1mM DSSO. After a dMP^78^ control of XL reaction that confirmed the XL-stabilization of both intact spliceosome and sub-complexes (**Supplementary Fig. 17c**), samples were digested and cleaned-up as described in our general workflow overview (**Fig. 1**). Using the 11-CV FAIMS-GPF protocol, we could identified 231, 283 and 335 unique XL in replicates 1, 2 and 3, respectively, for a total of 405 XLs, *i.e.* a 4 times increase compared to the analysis without FAIMS (100 XLs in total; **Supplementary Fig. 17d**). Among them, we validated 283 unique XLs (file threshold 2/3, 4.6 times increase, 72 % reproducibility) accounting for 91 PPIs including 110 proteins (**Fig. 5a**). We could map 55 XLs on the B complex structure (PDB: 8QO9), for which 93 % of XLs were distance-validated (**Fig. 5b, Supplementary Fig. 18**). The cross-links that could not be distance-validated may be due to the co-existence of other intermediates, such as pre-B complex in the sample (**Supplementary Fig. 19**). We also identified several PPIs that are likely involved in interactions mediated by fragments of the B complex components that were not present in the cryo-EM density. Closer inspection of the putative position of such fragments further supports the presence of an additional 21 intermolecular crosslinks on the B complex. Others detected PPIs occur between partners known to enter the splicing cycle later in the activation process (*e.g.* NineTeen Complex (NTC); also known as Prp19C)^79,80^ suggesting the presence of more activation-advanced spliceosome intermediates in this sample. Altogether, our fast and sensitive multiple-CV FAIMS-GPF XL-MS workflow is particularly well-suited for characterizing PPIs in complex spliceosome samples, enabling the detection of transient PPIs as well as providing a powerful tool to study sample heterogeneity.

## DISCUSSION

All cellular processes depend on interactions between biomolecules, particularly proteins. Thus, hundreds of thousands of PPIs drive most aspects of cellular function, and deepening our ability to identify and study PPIs is essential for advancing biological research. Therefore, developing new methods for detecting and identifying PPIs, especially transient PPIs formed by low abundance proteins remains important. In general, chemical crosslinking strategies are able to capture PPIs in the native environment and preserve physiologically relevant interactions, including those involving low affinity complexes and low abundance proteins. However, analyzing complex samples (such as cell lysates) that contain a mix of cross-linked and non cross-linked peptides, with cross-linked peptide typically present in much lower concentrations, remains a major challenge and require large amounts of starting material (> 500 µg), extensive in-solution fractionation (> 20 fractions), and increased instrument acquisition times (days to weeks). This can make XL-MS experiments impractical for samples like isolated organelles, nuclear extracts, specific cell lines or patient samples that are not available in such quantities.

Here, we addressed these challenges by developing an optimized XL-MS workflow based on FAIMS-GPF that reduces sample complexity at the MS level. This method represents a compromise between the amount of starting material required, the MS acquisition time, and the level of PPI detection. Overall, in all three case studies described above (purified trimeric CAK complex, HeLa cell lysate, and spliceosome), our optimized FAIMS-GPF XL-MS workflow resulted in increased sensitivity while starting with a few µg of sample and completing the triplicate experiments within 1 to 3 days. More specifically, applying 4-CV FAIMS-GPF on XL-MS CAK complex (1.6 µg per replicate) allowed deepening our understanding of PPIs by almost doubling the number of XL-IDs. At a proteome-wide level, the 11-CV FAIMS-GPF protocol using a total of 20 µg of starting material was able to provide 1269 XLs and 213 PPIs in less than 3 days, numbers that were previously only reached after extensive in-solution fractionation (*e.g.* using ≥ 20 strong cation exchange fractions for a total of ∼500 µg of starting material and requiring weeks of sample prep and instrument time)^27,28,62,63,81^. Therefore, our method represents an alternative to classical XL-MS workflows and provides a dramatic decrease of the amount of sample and instrument time needed, while retaining high sensitivity and reproducibility.

The improved sensitivity allowed us to make several important discoveries. For example, our analysis of the CAK complex led to the discovery that the N-terminal part of MAT1, mainly in the helical bundle, is able to fold back on the Cdk7/Cyclin H dimer. This interface mediated by MAT1 N-terminal was not observed in reported cryo-EM structures of CAK because these structures were not able to capture this region of MAT1 given its high flexibility^47,48^. Although the interface and its functional role(s) remain to be analyzed and validated via orthogonal methods in future studies, these results indicated the effectiveness of our FAIMS-GPF XL-MS workflow to capture transient interactions involving flexible domains.

Additionally, by analyzing the interactome of HeLa cell lysates we were able to identify 118 PPIs that are not documented in STRING database^49^. Of note, among the newly discovered PPIs, we could identify potentially novel partners of ENO1 and of different heat-shock proteins, demonstrating the potential of our approach to uncover biologically relevant interactions.

We also benchmarked the improved speed and sensitivity of FAIMS-GPF for the XL-MS analysis of challenging samples, such as the spliceosome. Spliceosome is a large macromolecular assembly composed of hundreds of proteins and RNAs that is often purified in low quantities, and with high heterogeneity. Here, we preceeded 11-CV FAIMS-GPF XL-MS with MP-based quality control steps of very high sensitivity. Using our protocol, we could perform a XL-MS experiment in triplicates from less than 40 µg of starting material within one week, which is almost impossible with more conventional XL-MS protocols. This is important because these amounts of starting materials (less than 40 µg) make the use of XL-MS fully compatible with modern structural biology (e.g. cryo-EM) workflows^74,75^.

Altogether, our results pinpoint the efficiency and versatility of FAIMS-GPF XL-MS workflows. This versatile and easily tunable method uses low sample amounts and reduces instrument time, while ensuring compatibility with a broad range of samples. We thus expect the FAIMS-GPF method will facilitate a more straightforward integration of XL-MS workflows with structural biology efforts aimed at analyzing complex and dynamic systems (such as the spliceosome), and in-routine with proteomics platforms. Ultimately, this will further boost our ability to characterize roles of protein-protein association networks interconnection in biological and cellular processes.

## MATERIALS AND METHODS

### Sample preparation and cross-linking reactions of CAK complex

For production of the CAK ternary complex, the genes coding for the CDK7 kinase fused to a C-terminal Strep-tag II sequence, cyclin H, and MAT1 were cloned in pAC8_MF as described in ^82^. Viruses were generated by co-transfection of the transfer vectors with the AcMNPV BAC10:KO1629, Δv-cath/chiA, mCherry bacmid and amplified using standard procedures^83^. Sf21 cells grown in suspension were infected with the corresponding virus, harvested after 3 days incubation at 27 °C, washed in PBS containing 10% v/v glycerol and flash frozen in liquid nitrogen. Typically, cells are grown in 2 L flasks containing 500 ml of medium and are infected at a density of 1.10^6^ cells/ml using 20 ml of P1 virus stock (assuming a titer of 5 × 10^7^ pfu/ml, it corresponds to a multiplicity of infection of 2).

For purification, the cell pellet from an infected suspension culture is disrupted by sonication in 20 mM Hepes-KOH pH8, 250 mM KCl, 0.1% Nonidet P40, 1 mM DTT and EDTA free protease inhibitor cocktail (Roche). After centrifugation at 25 000 g for 20 minutes, the clarifed lysate was incubated with Strep-Tactin®XT afnity chromatography (IBA-Lifesciences, 2-4030-010). After extensive washing with 20mM Hepes pH8, 150mM KCl, 1mM EDTA, 1mM DTT, bound proteins were eluted in the same buffer supplemented with 50 mM biotin. The CAK complex at 0.7 mg/mL in its native buffer was cross-linked in three independent replicates. 20 µg per replicates were incubated with DSSO freshly diluted in anhydrous DMSO to reach 200 molar excesses (45 min, RT). Reaction was quenched with 20 mM Tris HCl for 20 min.

CAK complex at 0.7 mg/mL in its native buffer (20mM Hepes pH 8, 150mM KCl, 1mM EDTA, 1mM DTT, 50mM Biotin) was cross-linked in three independant replicates. 20 µg per replicates were incubated with DSSO freshly diluted in anhydrous DMSO to reach 200 molar excesses (45 min, RT). Reaction was quenched with 20 mM Tris HCl for 20 min.

### Sample preparation and cross-linking reactions of spliceosome

The spliceosome sample was obtained by assembling spliceosomes from a nuclear extract previously treated 10 min at 30°C with 1 μM cyklin-dependant kinase inhibitor OTS964^84^ onto an AdML-3xMS2 substrate for 1h at 30°C essentially as described previously^65^. Assembled spliceosomes were purified by glycerol cushion, amylose affinity and glycerol gradient ultracentrifugation. In summary, 60 mL of splicing reaction was diluted by addition of 60 mL of dilution buffer K150 (20 mM HEPES-KOH pH 7.9, 40 mM KCl, 0.25 mM EDTA, 1% glycerol, 1 mM DTT, 0.03% NP40). The resulting diluted splicing reactions were loaded onto 9 ml of 40% glycerol cushion and ultra-centrifuged at 32,000rpm in SW32 Beckmann (20 mM HEPES-KOH pH 7.9, 50 mM KCl, 0.25 mM EDTA, 40% glycerol, 1 mM DTT, 0.02% NP40). Cushion were then recovered, diluted in 3,0 mL of D150K (20 mM HEPES-KOH pH 7.9, 150 mM KCl, 0.25 mM EDTA, 1% glycerol, 1 mM DTT, 0.03% NP40) and incubated with amylose resin high flow (NEB) overnight at 4°C. After extensive washing with WK150 (20 mM HEPES-KOH pH 7.9, 150 mM KCl, 0.25 mM EDTA, 1% glycerol, 1 mM DTT, 0.01% NP40) the MS2-containning complexes were eluted using WK150 supplemented with 12 mM of maltose. The fractions containing the complexes, as judged by fluorescein signal, were pooled, precipitated with PEG20K, resuspended in buffer W150K and loaded onto 10-30 % glycerol gradients (20 mM HEPES-KOH pH 7.9, 150 mM KCl, 0.25 mM EDTA, 10% or 30% glycerol, 1 mM DTT, 0.01% NP40) and spun for 3h at 50000 rpm in SW60 rotor, gradients were fractionated from the bottom to the top and the fluorescent signal measured and plotted against the number of fractions. The B-like complex peak fractions were pulled together and concentrated in 100 KDa to reach the concentration of roughly 1 mg/ml (500 nM).

Three 11 µg aliquots of spliceosome were independently cross-linked with 1 mM DSSO (45 min, RT), and quenched with 20 mM Tris HCl for 20 min.

### Native mass spectrometry measurements of CAK complex

CAK trimer was buffer-exchanged against 150 mM ammonium acetate using Zeba Spin desalting columns (Thermo Fisher Scientific, Rockford, IL, USA). Sample was analyzed in positive ion mode on an Orbitrap Exactive Plus EMR (Thermo Fisher Scientific, Bremen, Germany) coupled to an nano-electrospray source (TriVersa NanoMate, Advion Biosciences, Ithaca, U.S.A.), piloted by Xcalibur. Following parameters were used: resolution of 17500, in-source collision-induced dissociation (CID) of 150 eV, Collision Energy of 70 eV, trapping pressure of 7 (u.a.). The injection-, inter- and bent-flatapoles were set to 4V. Data were analyzed using Xcalibur Qual Browser and UniDec.

### Native and denaturing mass photometry measurements of CAK and spliceosome complexes

MP measurements were performed with a TWO^MP^ (Refeyn Ltd, Oxford, UK) at room temperature (18 °C). Microscope slides (24×50 mm, 170±5 µm, No. 1.5H, Paul Marienfeld GmbH & Co. KG, Germany) were cleaned with milli-Q water, isopropanol, milli-Q water and dried with a clean nitrogen stream. Six-well reusable silicone gaskets (CultureWell^TM^, 50-3 mm DIA x 1mm Depth, 3-10 µL, Grace Bio-Labs, Inc., Oregon, USA) were carefully cut and assembled on the cover slide center. After being placed in the mass photometer and before each acquisition, an 18 µL droplet of PBS was put in a well to enable focusing on the glass surface.

For native MP, samples were analyzed at 50 nM in a PBS droplet. Denaturing MP experiments were carried out as described previously^42^. Denaturation was done as described previously^78^, by incubating samples in 5.4 M Urea (Sigma, Saint-Louis, USA) for 5 min at 90 °C. Samples were drop-diluted to 50 nM and measured in a PBS droplet. Three movies of 3000 frames were recorded (60 s) for each sample using the AcquireMP software (Refeyn Ltd, Oxford, UK). A contrast-to-mass calibration was performed twice a day by measuring a mix of Bovine Serum Albumin (66 kDa), Bevacizumab (149 kDa), and Glutamate Dehydrogenase (318 kDa) in PBS buffer, pH 7.4. Data were finally processed using the DiscoverMP software (Refeyn Ltd, Oxford, UK).

### Sample preparation and cross-linking reactions of HeLa Lysate

For ‘native’ cell lysis, live HeLa cells were washed with PBS, pelleted with ultracentrifugation and frozen at - 80°C overnight. Pellet was unfrozen on ice in 500 µL of a non-denaturing buffer containing 20 mM Hepes, 150 mM NaCl, 1.5 mM MgCl2 and 0.5 mM DTT, adjusted at pH 7.8. A tablet of EDTA-free protease inhibitor cocktail was added per 10 mL of buffer. Cells were mechanically lysed on ice with 40 quick pushes using a 27 ¾ Gauge syringe. Cell debris were further removed using a 13,800 G centrifugation for 10 min at 4°C. Supernatant protein concentration was measured at 1.4 mg/mL using Pierce assay. Concentration was adjusted at 1 mg/mL and 3 aliquots of 100 µg of lysate were cross-linked with final concentration of 2 mM of DSSO (45 min, RT), followed by a quenching step with 20 mM TrisHCl. For FAIMS-GPF CV screening 50 µg of each XL replicate were pooled. For final FAIMS-GPF XL-MS experiments a 20 µg aliquot was taken for each independant XL replicate.

### SDS-PAGE separation of cross-linked samples

All cross-linked proteins and complexes were migrated to check the efficiency of XL reaction on in-house 12 % acrylamide denaturing SDS-PAGE gels (1.5 mm thickness). Volume corresponding to 1.5 µg of each XL sample (and non-XL controls) was diluted (1:1) with 2x concentrated Læmmli buffer (4 % SDS, 20 % glycerol, 10 % 2-mercaptoethanol, 0.01 % bromphenol blue and 0.125 M Tris HCl) and incubated 5 min at 95°C. After sample loading, gels were migrated at 50 V for 20 min, 100 V until the 2/3 of the gel and 120 V until the end. After migration, gels were fixated for 20 min (3 % phosphoric acid, 50 % ethanol), washed 3×20min with milli-Q water and stained overnight with Coomasie Brillant Blue (G250, Sigma, Saint-Louis, USA). They were finally rinced 3×20 min with milli-Q water.

### Proteolysis of samples and automated cleanup of peptides

All samples were denatured (6M Urea), reduced (5 mM DTT, 30 min at 37 °C) and alkylated (15 mM Iodoacetamide, 1 hour in the dark, RT). After dilution to < 2M Urea using Ammonium Bicarbonate aliquots were digested overnight at 37°C with Trypsin/LysC (Promega, Madison, USA) at a 25:1 enzyme:protein (w/w) ratio for XL-lysate and 50:1 for CAK. Digestions were quenched with 1 % TFA. Peptides generated by proteolysis were cleaned up using the AssayMAP Bravo platform (Agilent Technologies; Santa Clara, California) with 5 μL C18 cartridges (Agilent). Briefly, cartridges were primed (100 μl 0.1 % TFA in 80 % CAN), equilibrated (50 μl 0.1% TFA in water), 180 μl of digested peptides diluted in equilibration buffer were loaded on the cartridges, and finally cleaned with 50 μl of equilibration buffer. Peptides were eluted with 50 µl 0.1 % TFA in 80 % ACN. Eluted peptides were frozen and kept at -80 °C until analysis.

### nLC-MS/MS proteomics experiments on HeLa lysates digests

Analysis were done using a Dionex UltiMate 3000 RSLC nano system (Thermo Scienctific) coupled to an Orbitrap Eclipse Tribrid (Thermo Scientific). We used a mobile phase A (0.1 % FA in water) and mobile phase B (0.1 % FA in 80 % ACN/20 % Water). 400 ng of peptides were injected and first trapped on an Acclaim PepMap 100 C18 20 mm x 0.1 mm, 5 µm diameter particles pre-column (Thermo Scientific) at 10 µL/min for 3 min. Separation was done at 300 nL/min on an Aurora Series C18 UHPLC column (250 mm x 75 µm, 1.6 µm diameter particles from IonOpticks) with the following gradient: 2.5 % B for 3 min, 32.2 % B at 53 min, 50 % B at 63 min, 98.0 % B at 65 min, 98.0 % B kept until 70 min, 2.5 % B at 71 min. Analysis were carried in Data Dependant Acquisition (DDA) with a 2s cycle, full MS spectra were acquired over a 300-1800 m/z range at a resolution of 120000 in the Orbitrap, with an standard AGC target and an Auto max. injection time. For MS/MS, intensity threshold was set at 5e3, and 2-7+ charges were selected for Higher Collisional Dissociation (HCD) using respectively 30 % Normalized Collision Energy (NCE). MS/MS spectra were acquired in the Ion Trap with a Rapid scan rate and normal AGC target and an Auto max. injection time of 200 ms. MaxQuant was used to process proteomics experiments on lysate using standard parameters. To compare proteins abundances, mean of LFQ value was calculated between replicates, ignoring missing values.

### nLC-MS/MS of cross-linked samples without FAIMS

For nanoLC-MS/MS/ analysis, cross-linked peptides were dried in a SpeedVac concentrator and resuspended in 2% ACN/0.1% formic acid. Analysis were done using a Dionex UltiMate 3000 RSLC nano system (Thermo Scienctific) coupled to an Orbitrap Eclipse Tribrid (Thermo Scientific). We used a mobile phase A (0.1 FA in water) and mobile phase B (0.1 % FA in 80%ACN/20%Water). 400 ng of peptides were injected and first trapped on an Acclaim PepMap 100 C18 20 mm x 0.1 mm, 5 µm diameter particles precolumn (Thermo Scientific) at 10 µL/min for 3 min. Separation was done at 300 nL/min on an Aurora Series C18 UHPLC column (250 mm x 75 µm, 1.6 µm diameter particles from IonOpticks) with the following gradient for CAK: 2.5 % B for 3 min, 43.8 % B at 93 min, 98.0 % B at 95 min, 98.0 % B kept until 100 min, 2.5 % B at 102 min. Following gradient was used for XL-lysates: 2.5 % B for 3 min, 43.8 % B at 123 min, 98.0 % B at 125 min, 98.0 % B kept until 130 min, 2.5 % B at 132 min. Analysis were carried in Top10 Data Dependant Acquisition (DDA), full MS spectra were acquired over a 300-1800 m/z range at a resolution of 120000 in the Orbitrap, with an AGC target of 750 % and a 100 ms max. injection time. For MS/MS, intensity threshold was set at 2.0e5, and only 3-7+ charges were selected for Higher Collisional Dissociation (HCD). HCD was applied in Stepped Collision Energy mode using respectively 21, 26, 31 % Normalized Collision Energy (NCE). MS/MS spectra were acquired at 60000 of resolution (1.6 m/z isolation window) in the Orbitrap using an AGC target of 400 % and a max. injection time of 200 ms.

### nLC-FAIMS-MS/MS gas-phase fractionation experiments for cross-linked samples

For all samples, nanoLC separation 90 min separation/105 min gradient was used (see above for CAK complex). MS analysis were carried in Top10 DDA, with full MS spectra acquired over a 300-1800 m/z range at a resolution of 120000 in the orbitrap, with an AGC target of 250 % and a 100 ms max. injection time. For MS/MS, intensity threshold was set at 2.5e4, and 3+ to 7+ charge states were selected for Higher Collisional Dissociation (HCD). HCD was applied in Stepped Collision Energy mode using 21, 26, 31 % Normalized Collision Energy (NCE). MS/MS spectra were acquired at 60000 of resolution (1.6 m/z isolation window) in the Orbitrap using a standard AGC target, an auto max. injection time and a dynamic exclusion time of 60 s. For FAIMS setting, gas mode was set as static with a total carrier gas flow of 4.6 L/min and a standard FAIMS resolution. A single Compensation Voltage was use per each nLC-FAIMS-MS/MS run. For CV screening CV values used were: -40 V, -45 V, -50 V, -55 V, -60 V, -65 V, -70 V, -75 V, -80 V, -85 V, -90 V. For 4-CV GPF analysis of XL-CAK, -55 V, -60 V, -65 V, and -70 V were used; for 11-CV GPF analysis of XL-lysates and spliceosome, all CVs from - 40 to – 90 V were used.

### Data processing

For data processing, .raw data were directly processed with Thermo Proteome Discoverer 2.5.0.400 (Thermo Scientific) using the Sequest HT node for the identification of peptides and the XlinkX node for identification of crosslinks. For CAK a database containing only proteins of interest was used, and for proteome-wide samples, database containing full human proteome was used (retrieved the 2023/07/09). Cysteine carbamidomethylation was set as fixed modification and Methionine oxidation, N-term acetylation, tris-quenched mono-links and water-quenched mono-links were set as dynamic modifications. Trypsin was set as the cleavage enzymes with minimal length of 7 amino acids, 2 (linear peptides) and 3 (cross-linked peptides) missed cleavages were allowed, respectively for proteomics and XL identifications. For CAK we considered K, S, T, Y as DSSO linkage sites, while only K was considered for spliceosome and proteome-wide samples. Mass accuracies for both XL and linear peptides search were set to 10 ppm for MS1 and 20 ppm for MS2. To increase confidence, identification were only accepted for Maximal XlinkX scores > 40 and Δ_score_ > 4. A 1% false discovery rate was applied for both linear and XL-peptides at CSM (separated FDR between intra and inter-XLs), at a peptides and proteins level. All FAIMS-GPF XL-MS experiments on CAK, HeLa lysate and Spliceosome, including CV screening, have been deposited to the ProteomeXchange Consortium via the PRIDE^85^ partner repository with the dataset identifier PXD064228.

GO enrichment analysis was done by analyzing cross-linked proteins only identified in proteome wide FAIMS-GPF experiments in Metascape online server^86^. XL-MS data were visualized using xiVIEW webserver (www.xiview.org)^21^, to produce circular interaction networks and represent XL sites on protein sequences. Finally, validated cross–links were plotted different structures using the PyMol Molecular Graphics System (version 2.5.4, Schrödinger, LLC), ChimeraX as well as xiVIEW server to visualize and measure XLs Cα-Cα distances on the structure. Corresponding distances were validated if validating a cutoff of 30 Å.

## DATA AVAILABILITY

Complete FAIMS-GPF XL-MS dataset including CV screening on CAK and HeLa lysate, as well as the triplicate experiments on CAK, HeLa and Spliceosome have been deposited to the ProteomeXchange Consortium via the PRIDE^85^ partner repository with the dataset identifier PXD064228. Raw data from nMP datasets will be made fully accessible upon request. dMP profiles have be included in ProteomeXchange submission.

## ACKNOWLEDGMENTS

Authors would like to acknowledge Sarah Delaux for native mass spectrometry on CAK. This work was supported by the CNRS, the University of Strasbourg through the labex program (ANR-10-LABX-0030-INRT under the program Investissements d’Avenir ANR-10-IDEX-0002-02), the Ligue nationale contre le cancer (Ligue), the “Agence Nationale de la Recherche” (ANR-23-CE44-0035, ANR-20-CE12-0044), the French Proteomics Infrastructure (ProFI; ANR-10-INBS-08-03) and the French Infrastructure for Integrated Structural Biology (FRISBI, ANR-10-INBS-0005) as part of Instruct-ERIC. H.G.F. acknowledges the French Ministry for Education and Research for funding of his PhD. The work in C.C.’s group was supported by the ERC StG SPLIFEM and IdEx *Attractivity* from University of Strasbourg.

## AUTHOR CONTRIBUTIONS

H.G.F performed the mass photometry and XL-MS experiments. H.G.F., S.C conceived the study. S.C supervised the work. H.G.F., S.T, C.C., A.P and S.C wrote the manuscript. S.T. produced the spliceosome sample. S.P produced the CAK samples. H.G.F. and A.P analyzed and interpreted the CAK crosslinks. S.T., H.G.F. and C.C. analyzed and interpreted spliceosome crosslinks. D.K. and C.B contributed reagents and technical assistance. All authors proofread the manuscript.

## COMPETING INTERESTS

The authors declare no competing financial interest.

## Notes

### Competing Interest Statement

The authors have declared no competing interest.

