## Supplementary Data for "FAIMS-GPF XL-MS: crosslinking-mass spectrometry based on gas-phase fractionation"

*Gizardin-Fredon et al.*

**SUPPLEMENTARY FIGURES**

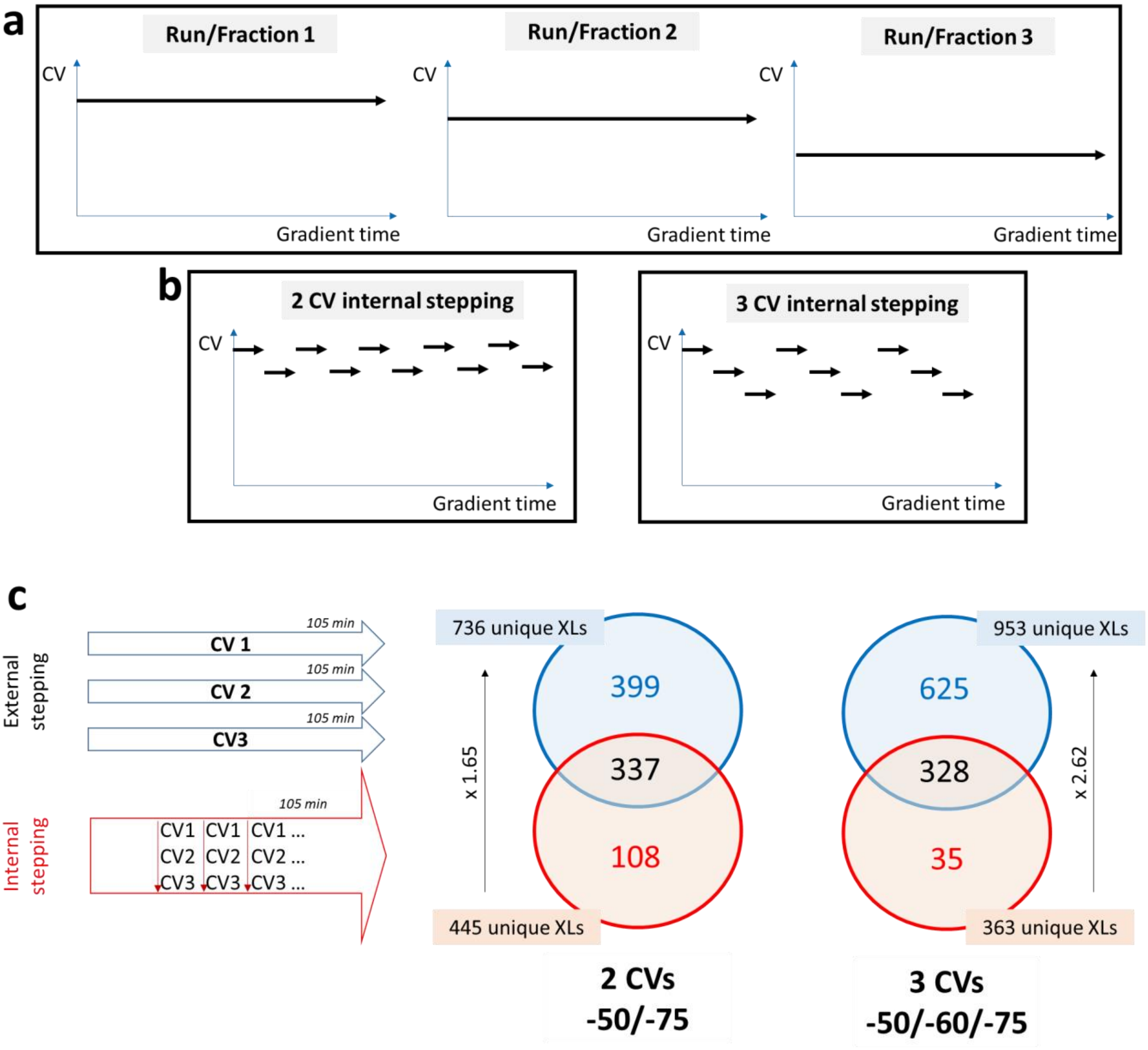


**Figure S1. Comparison of FAIMS external and internal stepping on an strong cation exchange (SCX)-enriched XL-Hela lysate (fraction number 6/6). (a)** Description of external stepping GPF experiments. **(b)** Description of internal stepping GPF experiments. **(c)** We first compared internal and external stepping for 2 (-50/-75V) and 3 (-50/-60/-75) CV values on a DSSO cross-linked HeLa lysate. We used a 105 min gradient and found that external stepping allowed 1.6 and 2.6 times more unique XLs identification for 2 and 3 CVs respectively, than internal stepping. As the sampling of precursors across gradients is poor with external stepping, many precursors will only be submitted to 1 CV value.

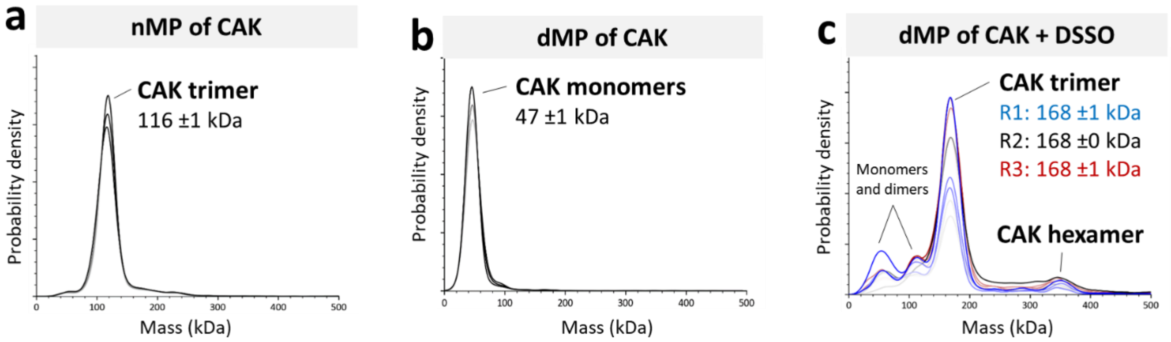


**Figure S2. Native and Denaturing MP experiments on XL-CAK complex. (a)** Native MP measurement of non-XL CAK, **(b)** dMP measurement of non-XL cak control, **(c)** dMP experiments of XL-CAK replicates, Mass shift between theory and XL-CAK measured mass in dMP corresponds to binding of mono-links and cross-links. SD comes from independent measurement (n=3).

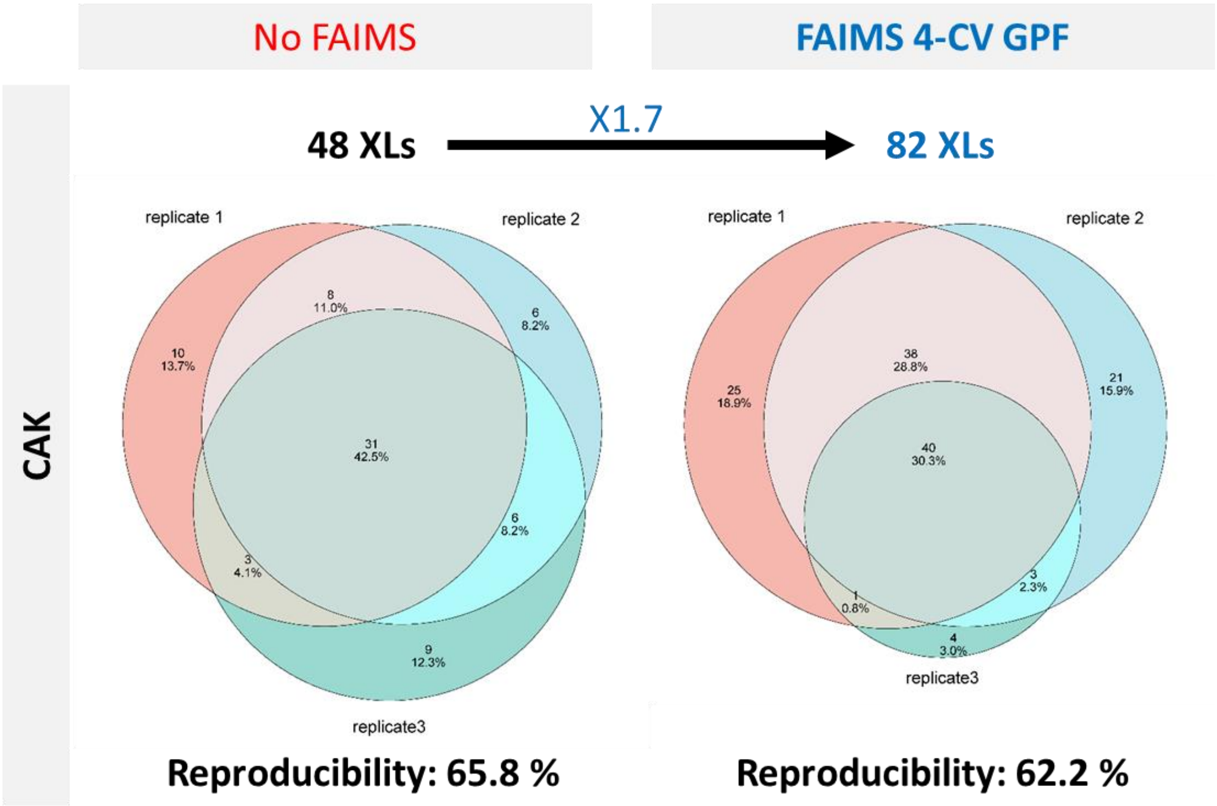


**Figure S3. Reproducibility between independant XL replicates for XL-MS experiments on CAK without and with FAIMS.**

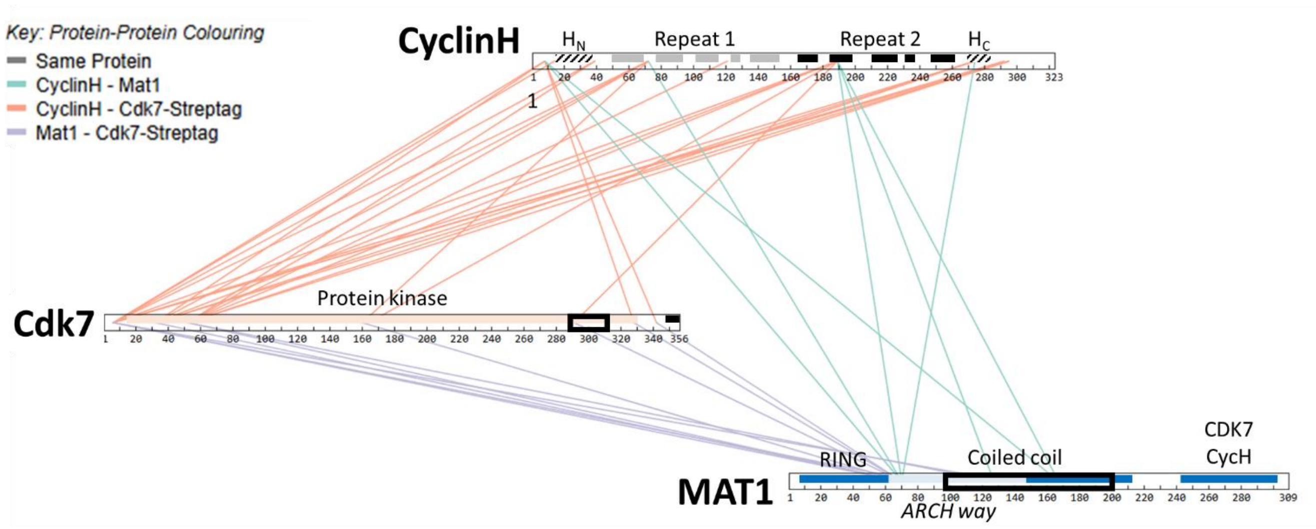


**Figure S4. Validated inter-protein XLs from the 4-CV GPF experiments mapped on the CAK partners sequences.** The different domains of protein partners are indicated. Regions of Cdk7 and MAT1 only covered using FAIMS-GPF are circled with black rectangles.

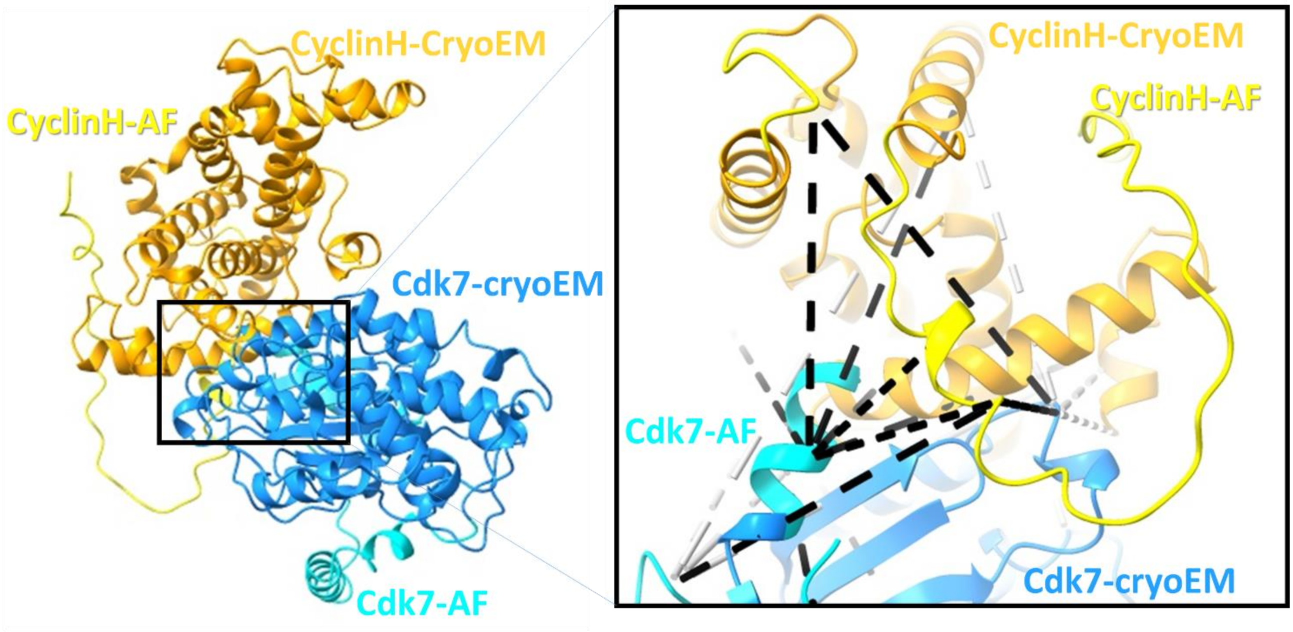


**Figure S5. Mapping of XLs of the CAK dataset on the structure of the model of the Cdk7/cyclin H binary complex (insert) obtained by superimposing the AF models of Cdk7 and Cyclin H on the cryo-EM structure of CAK (PDB: 8P6Y).** Residues modelled from the cryo-EM map are shown in blue (CdK7) and gold (cyclin H) while those that could not be located in the experimental map and are modelled from AF prediction are reprinted in cyan (Cdk7) and yellow (cyclin H). XLs between CAs of two residues are represented by grey (in the case of 2 amino acids modeled from experimental data) or back (when 1 or the 2 amino acids modelled from AF predictions).

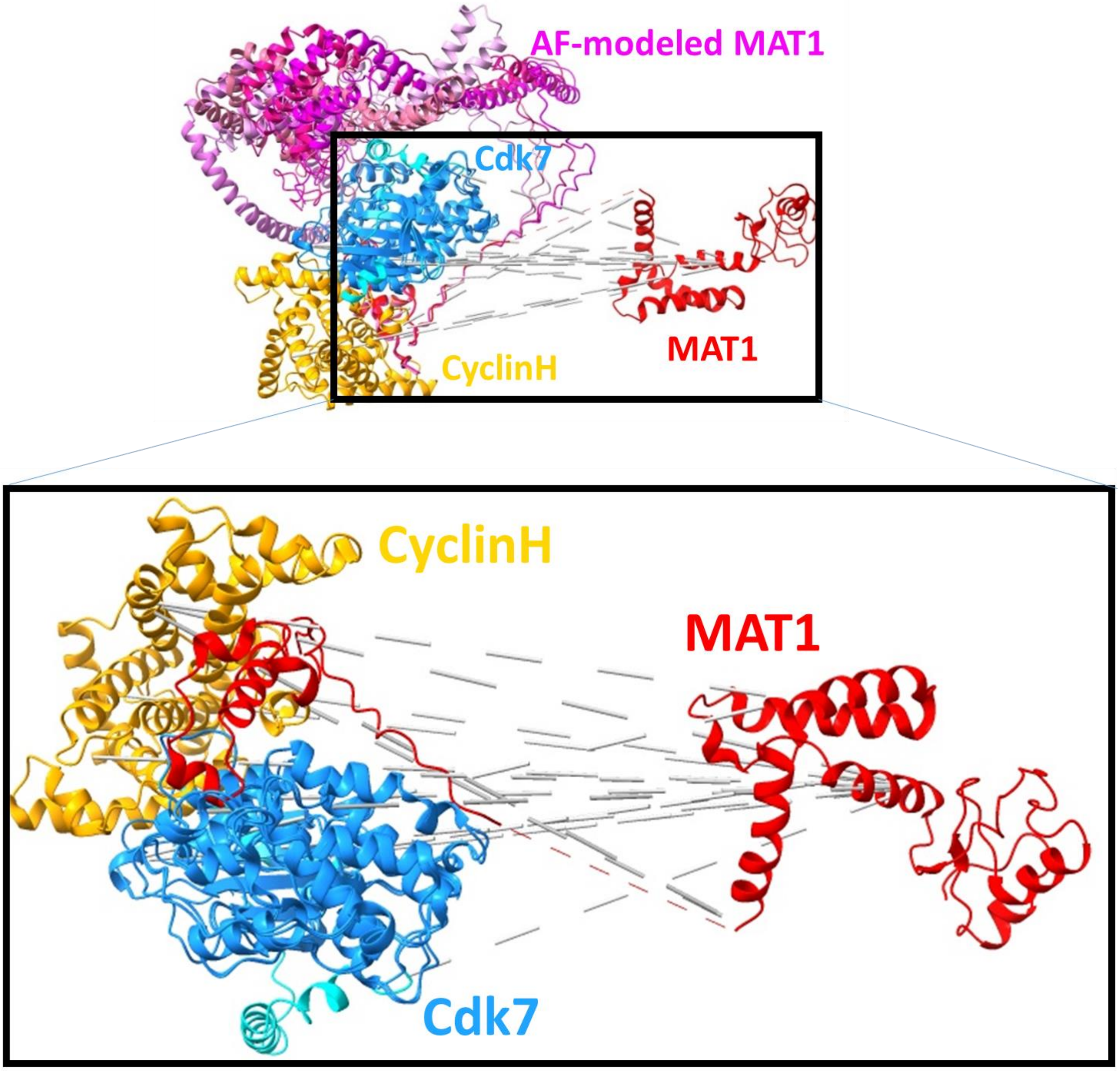


**Figure S6. Mapping of XLs of the CAK dataset on structure of CAK extracted from that human mediator–RNA polymerase II pre-initiation complex (PDB: 7NRV)**. Cdk7 is shown in blue, cyclin H in gold and MAT1 in red. XLs between are represented as dashed lines drawn between the Cα's of the two residues. In the insert the ribbon models MAT1 (in magenta) obtained of from 5 AF predictions of CAK MAT1 are superimposed.

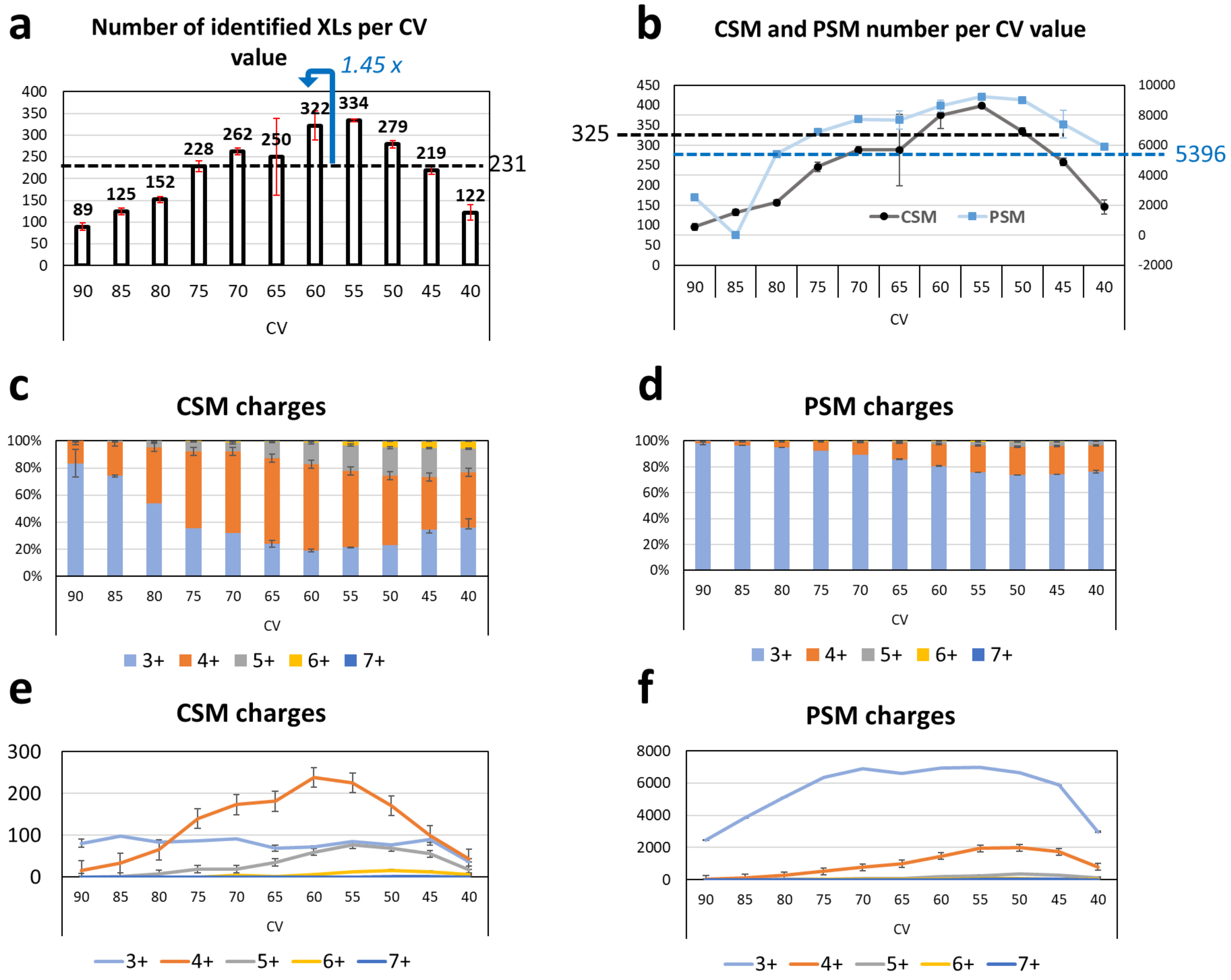


**Figure S7. Details of CV screening on XL-Hela lysate, using injection replicates (n=3). (a)** Number of unique XLs identified per CV value. All CSMs' SD are below 5 % apart from -65 V. (**b**) Number of CSM and PSM number per CV value identified with values obtained without FAIMS represented in dashed lines. (**c**) Detail of XLs charges states filtered, showed as proportion of CSM precursor charges for each CV value, (**d**) Detail of peptides charges states filtered, showed as proportion of PSM precursor charges for each CV value. (**e**) Detail of XLs charges states filtered, showed as absolute numbers of CSM precursor charges for each CV value. (**e**) Detail of peptides charges states filtered, showed as absolute numbers of PSM precursor charges for each CV value

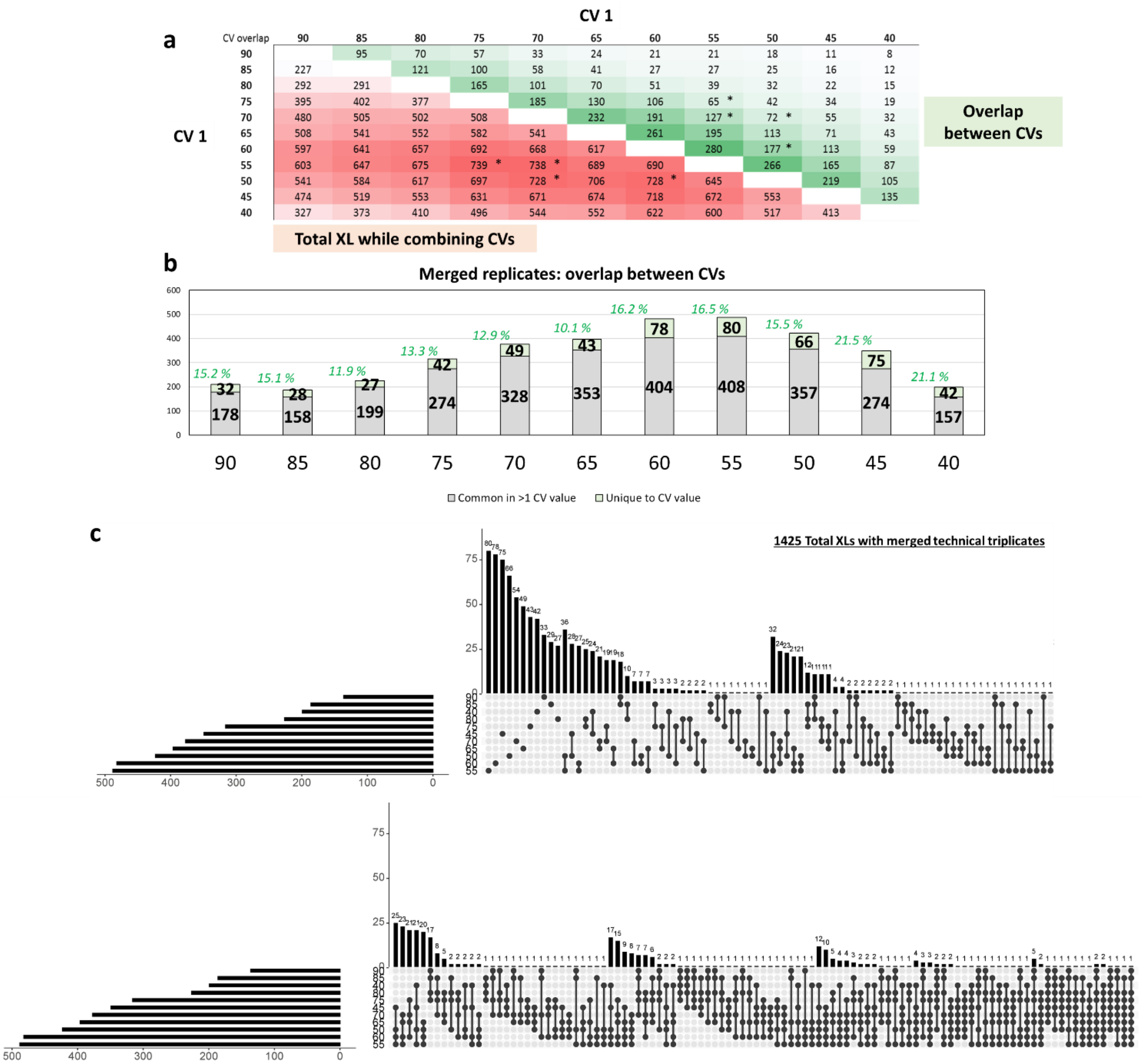


**Figure S8. Choice of best CV associations using merged technical triplicates. (a)** Matrix showing associations of CV pairs with in red number XLs identified when combining the 2 CV and in green number of XLs found in common in both CVs. Best combinations of CV pairs (maximizing red values and minimizing green values) are highlighted with stars. **(b)** Number of XLs identified per CV divided in XLs uniquely found to in the CV value as well as XLs present in at least CV value, from merged technical replicates (n=3)**. (c)** upsetR plot showing the overlap between all CV values**.**

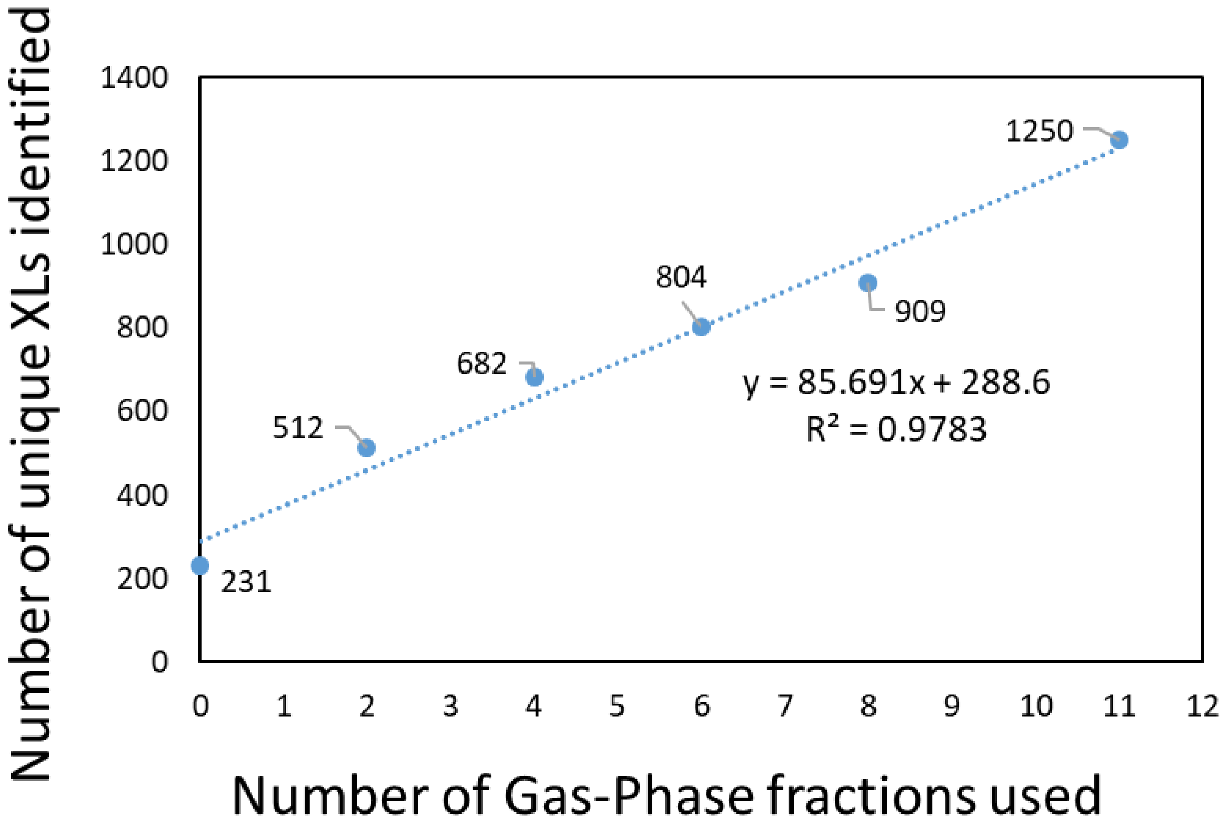


**Figure S9. Relation (linear fitting) between number of gas-phase fractions used and number of unique XLs identified for XL-HeLa lysate.**

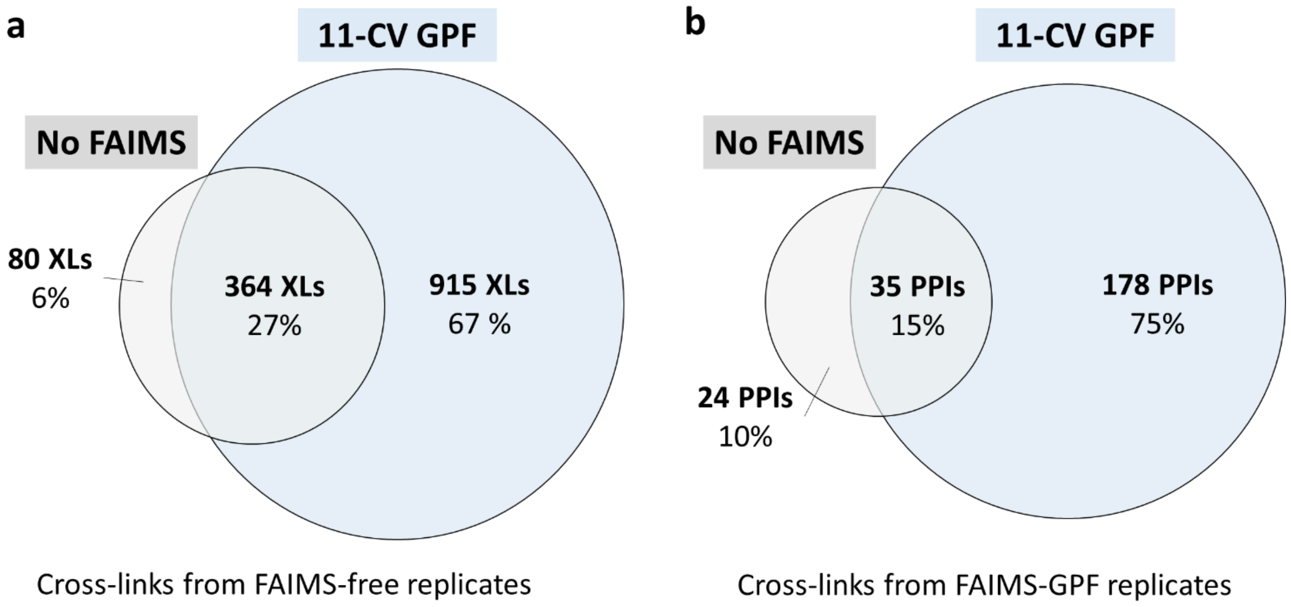


**Figure S10. Overlap between (a)** XLs identifications and **(b)** PPIs between FAIMS-free and 11-CV GPF experiments.

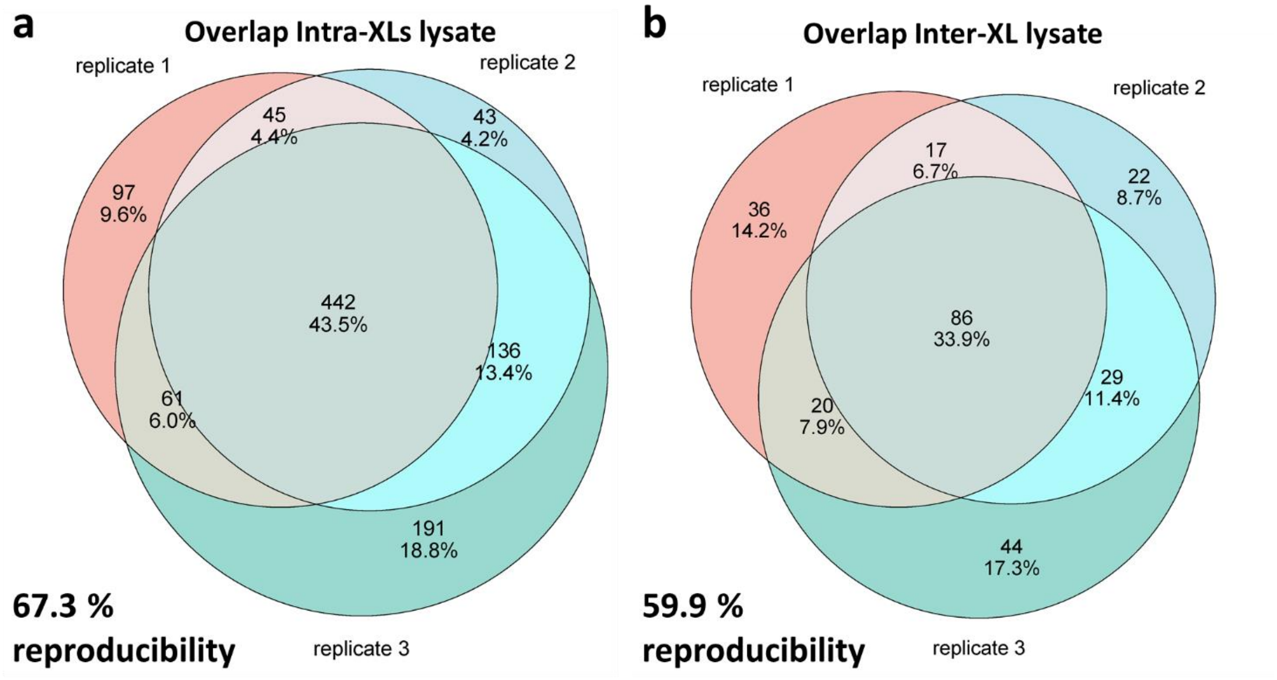


**Figure S11. Reproducibility of independent XL replicates of FAIMS-11-CV experiments at a proteome wide level for (a)** intra- and **(b)** inter-XLs. Overlap between FAIMS-free and 11-CV GPF experiments

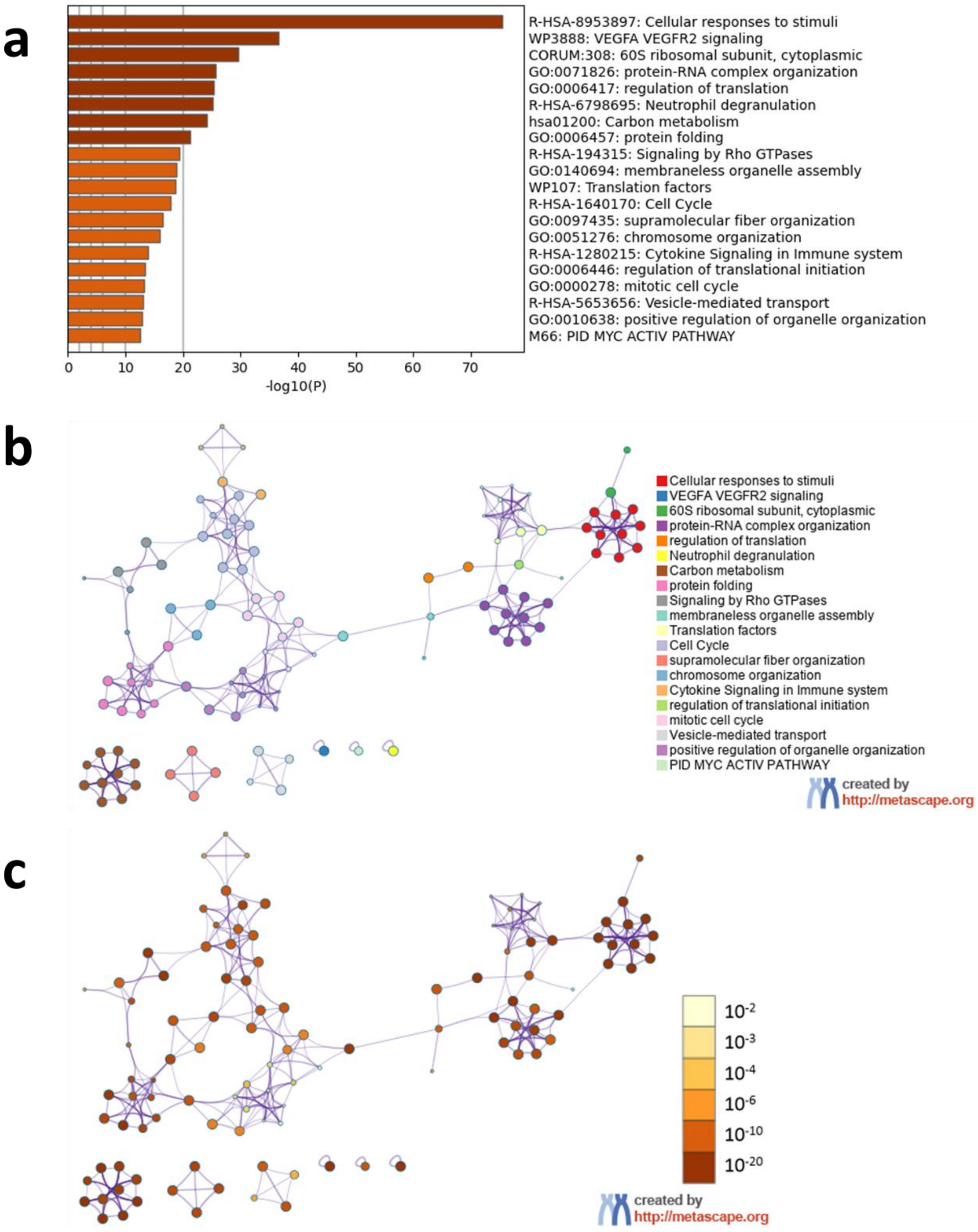


**Figure S12. Metascape[86] Gene Ontology enrichment analysis of cross-linked proteins only identified using FAIMS-GPF. (a)** Bar graph of enriched terms across input gene lists, colored by p-values**.** Network of enriched terms: **(b)** colored by cluster ID, where nodes that share the same cluster ID are typically close to each other; **(c)** colored by p-value, where terms containing more genes tend to have a more significant p-value**.**

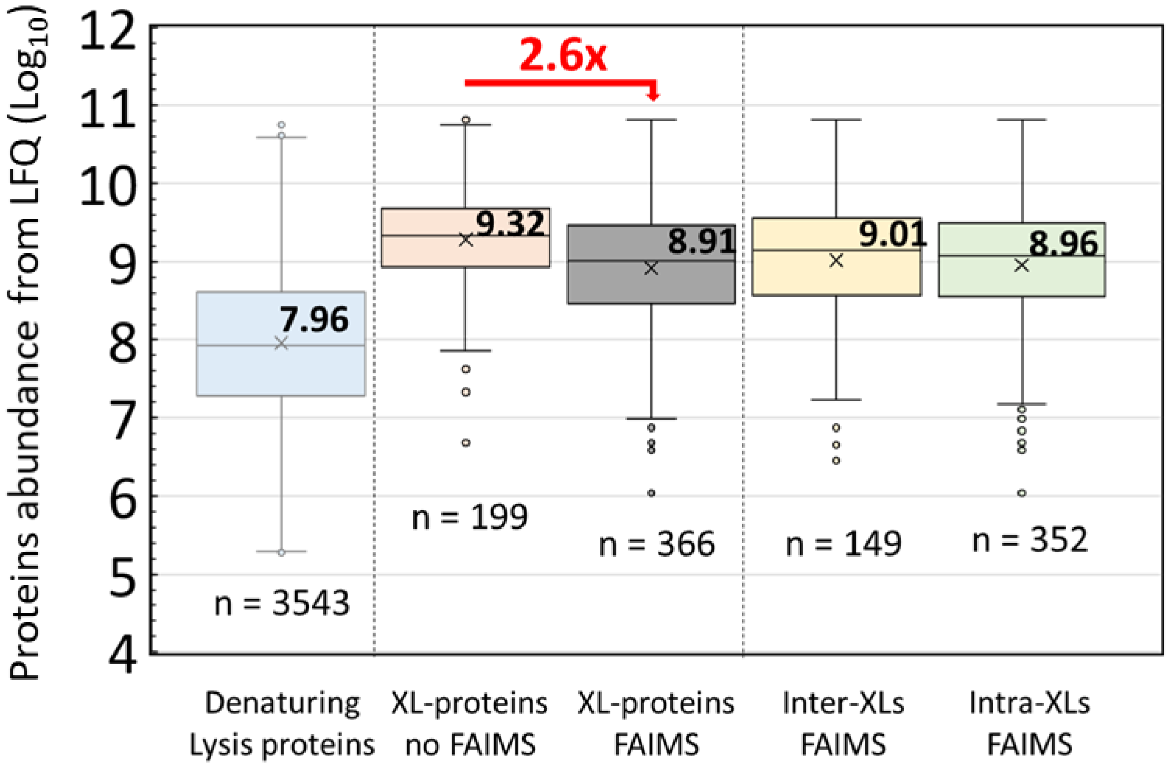


**Figure S13. Box-plot compares the LFQ protein abundance in whole cell lysate and abundances of cross-linked proteins.** Compared to the analysis without FAIMS, 11-CV GPF allows to reach proteins of 2.6 times less abundant. No major difference where found between abundance of protein with intra and inter-XLs. Values come from merged biological replicates and n displays number of values that could be plotted per category.

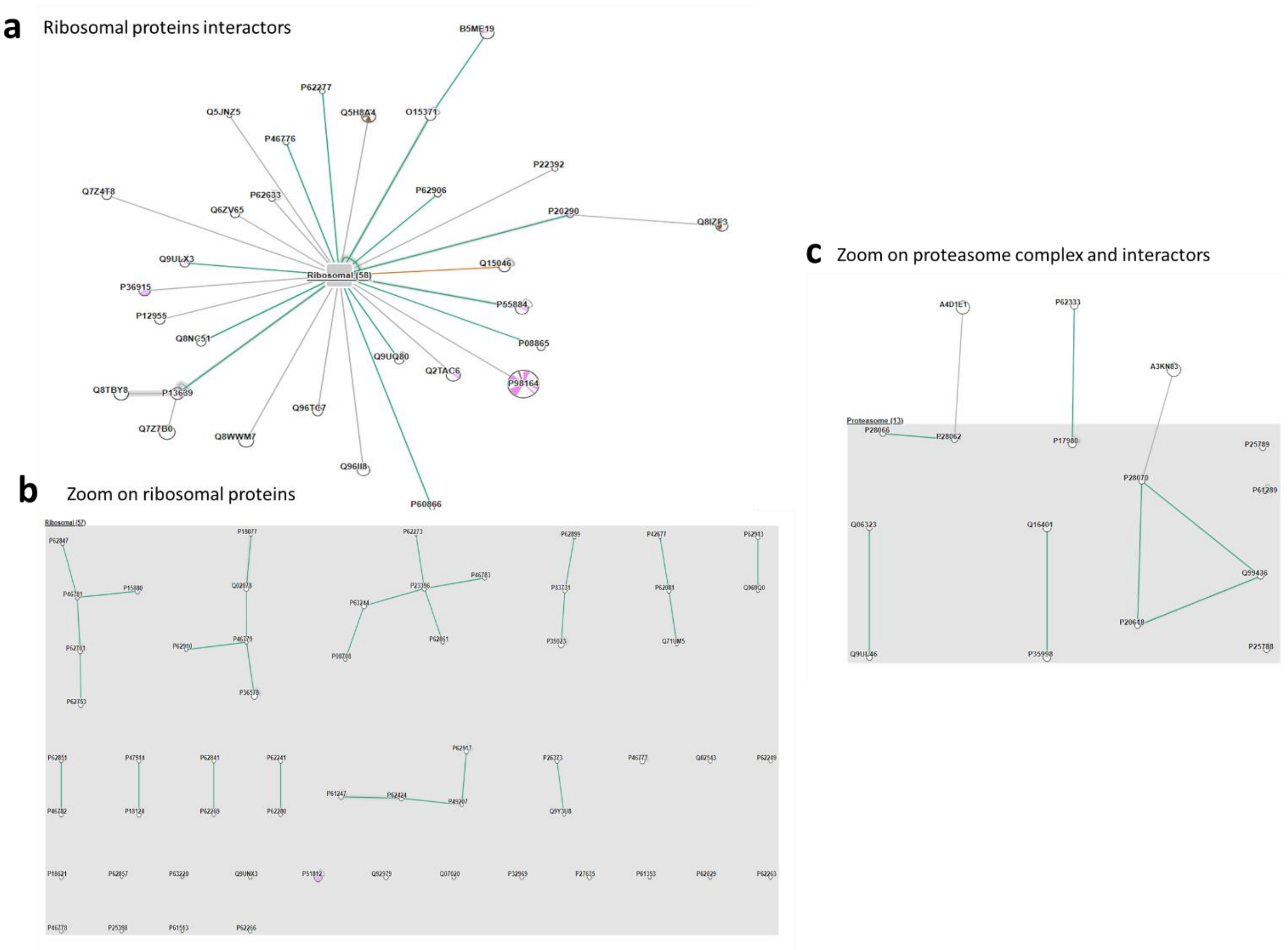


**Figure S14. (a)** Zoom on ribosomal interactors identified**. (b)** Zoom on inter-ribosomal proteins interactions, **(c)** Zoom on proteasome proteins interactions.

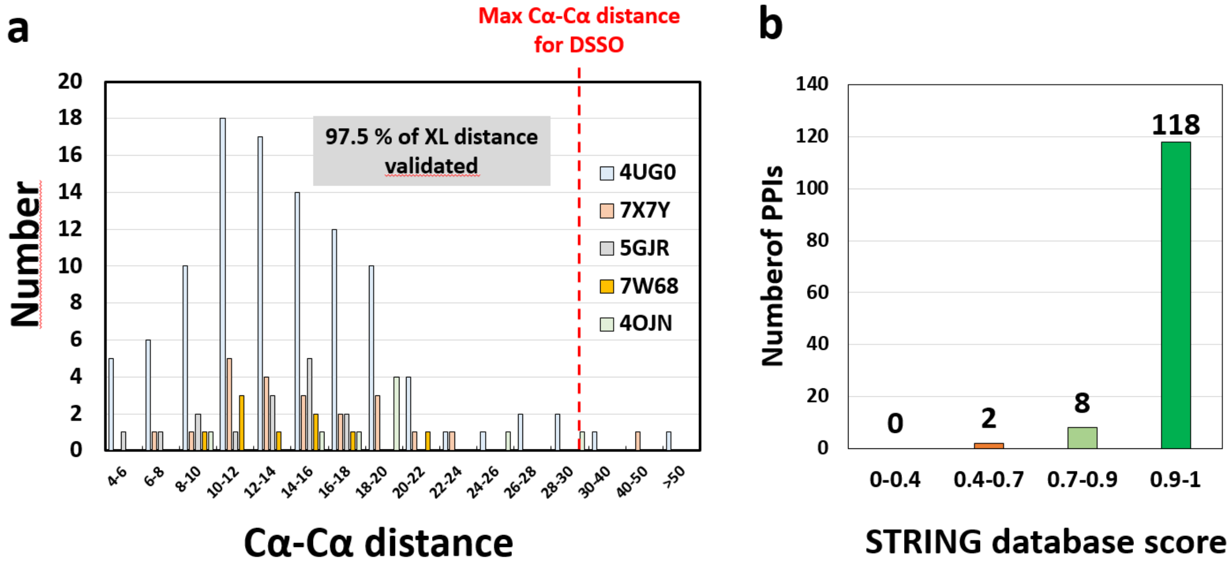


**Figure S15. Specificity of FAIMS 11-CV GPF dataset**. (**a**) Ca-Ca distances measured after mapping of intra- and inter-XLs on 5 known structures exhibit a high percentage of validation (97.5 %), (**b**) Scores of PPIs identified present in STRING database.

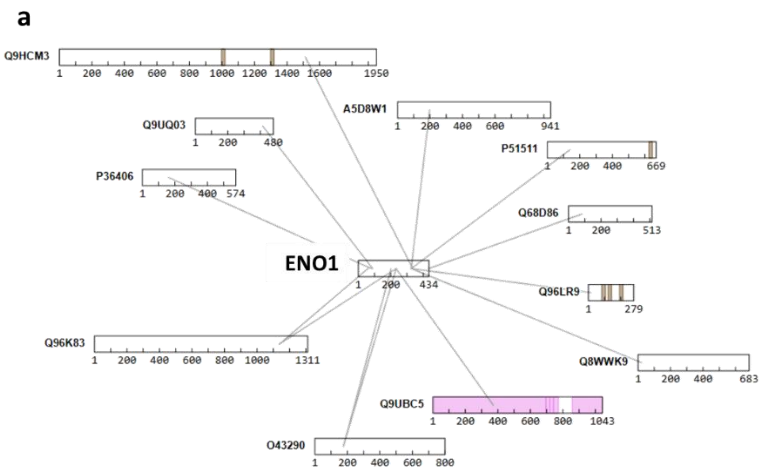


| Protein 1 | Protein 2 | Potential biological Implications |
|---|---|---|
| ENO1 | ZNF521 | May suggest a role in nuclear functions or transcriptional regulation. |
| ENO1 | CKAP2 | May imply involvement in microtubule-associated functions and cell cycle regulation. |
| ENO1 | Coronin-2B | Could reinforce ENO1's link to actin dynamics and cell motility. |
| ENO1 | SNUT1 | SNUT1 is a spliceosomal component. Interaction with ENO1 may suggest a nuclear role in RNA processing. |
| ENO1 | MYO1A | This could hint at ENO1's involvement in membrane remodeling or vesicular transport. |
| ENO1 | PLD1 | May link ENO1 to signaling cascades or membrane-associated functions. |
| ENO1 | KIAA1549 | Could represent an unexplored function or tissue-specific interaction. |
| ENO1 | C10orf2B | Might indicate a novel or tissue-specific function. |

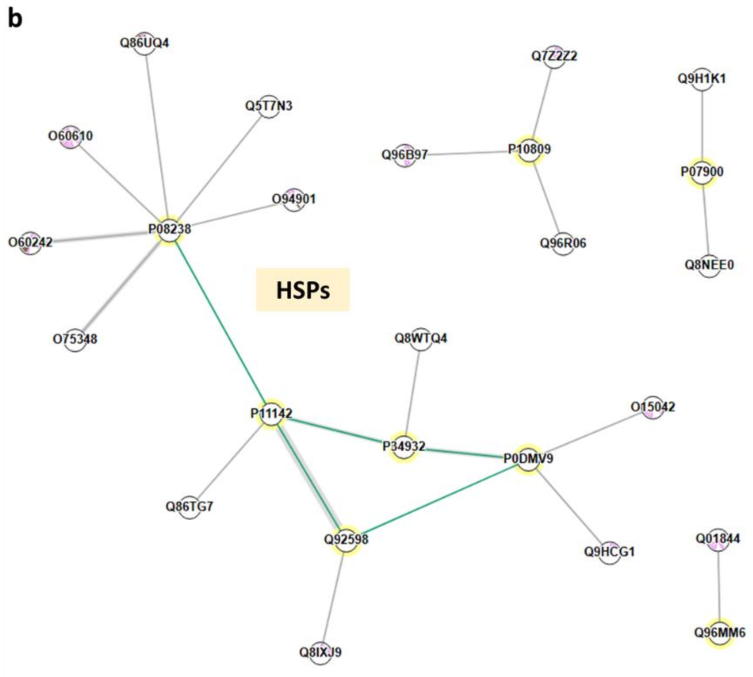


| Protein 1 | Protein 2 | Potential biological Implications |
|---|---|---|
| HSP90AB1 | DNAJB1 | Suggests crosstalk between Hsp40 co-chaperones and Hsp90 chaperone systems. |
| HSPA1B | U2SURP | May suggest regulation of RNA splicing via Hsp70 interactions. |
| HSP90AB1 | DNAJC7 | DNAJC7 links Hsp70 and Hsp90 systems; suggests multi-chaperone assembly. |
| HSP90AB1 | DIAPH1 | Suggests role of Hsp90 in cytoskeletal regulation. |
| HSP90AA1 | DNAJC3 | May link Hsp90 to ER stress pathways. |
| HSPB1 | DNAJB1 | Possible collaboration between small HSP and Hsp40 in stress response. |
| HSPA1B | HSPA4 | May represent functional interaction between Hsp70 isoforms. |
| HSPA4 | DNAJC10 | May suggest ER-based folding roles. |
| HSP90AB1 | DNAJC3 | ER-related co-chaperone possibly linking to Hsp90 pathway. |
| HSP90AB1 | DNAJC13 | May implicate Hsp90 in vesicle trafficking processes. |

**Figure S16. Interaction networks of (a) ENO1 and (b) Heat-shock proteins**. Interactions reported in STRING database are represented in green, known protein domains in Uniprot are highlighted in pink, transmembrane regions are highlighted in brown. Tables describe unreported interactions and their potential biological implications.

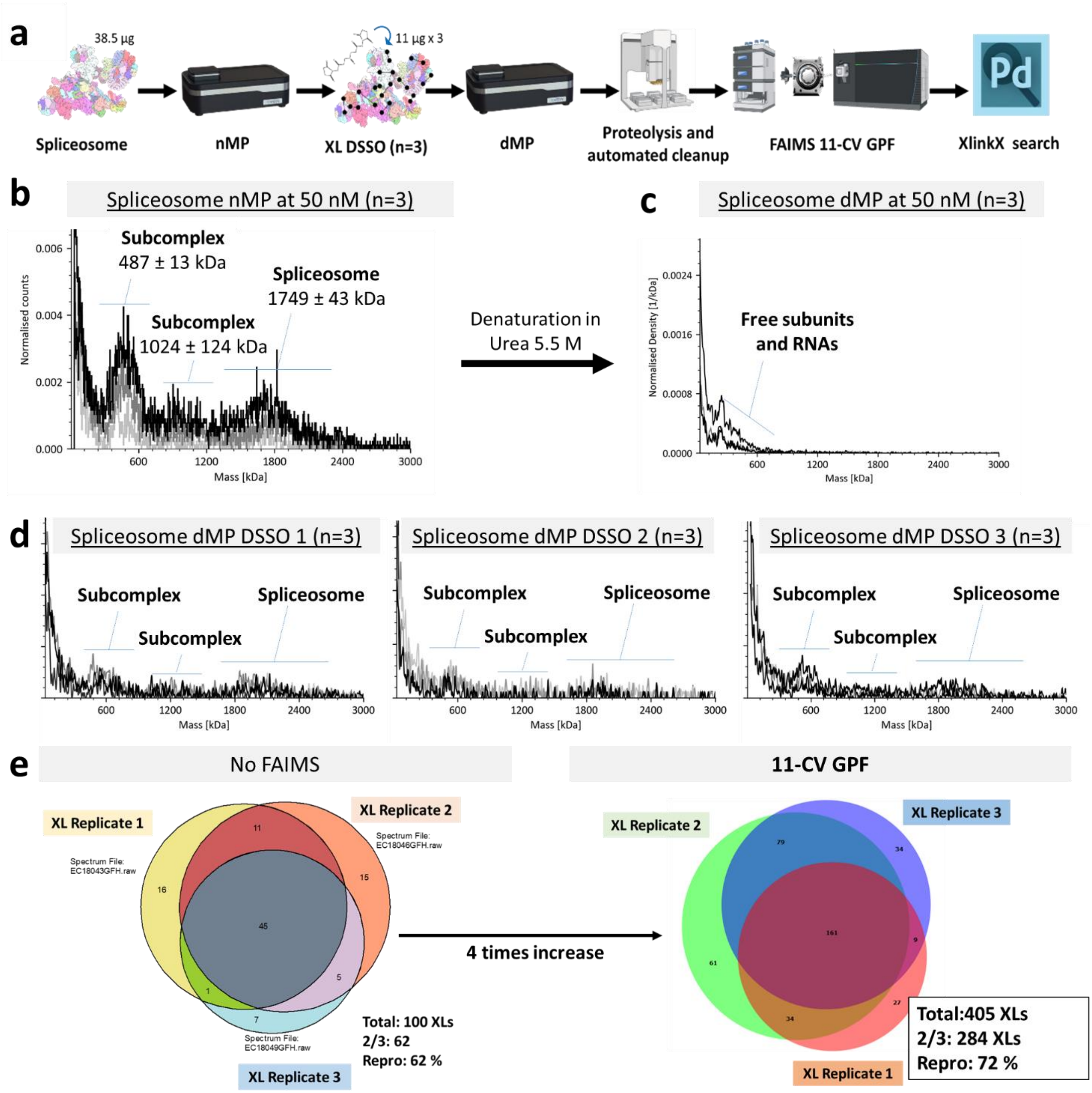


**Figure S17. Description and validation of spliceosome XL-MS dataset. (a)** Worfkflow used for the FAIMS-based XL-MS triplicate experiments on spliceosome. **(b)** nMp analysis of intact spliceosome highlights the co-existence of two subcomplexes. **(c)** dMP control of non-XL spliceosome shows the complete denaturation into RNA and protein subunits. **(d)** dMP control of XL reaction replicates highlights the presence of XL-stabilized sub-complexes and whol spliceosome in each replicates. **(e)** On the left, venn diagram shows the overlap between unique XLs identified in XL replicates analyzed without FAIMS. On the right, venn diagram shows the overlap between unique XLs identified in XL replicates analyzed using 11-CV FAIMS GPF. Reproducibility is increased by 10 % when using FAIMS-GPF and number of all XLs as well as reproducible XLs is multiplied by >4.

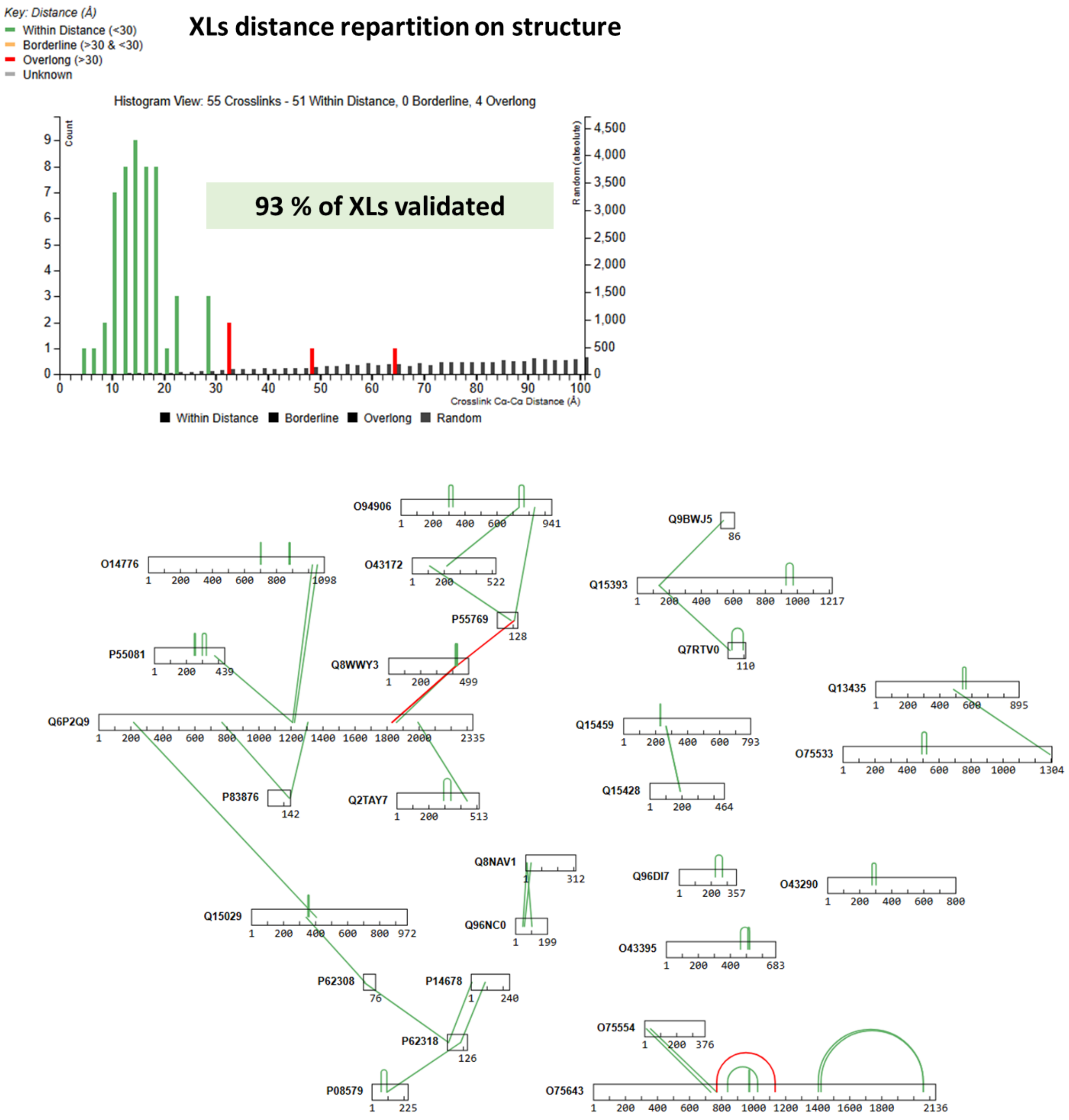


**Figure S18. Details of XLs mapping on B-complex structure.** On the top, XLs distance repartition. On the bottom, network of XLs that could be mapped on 8QO9 PDB structure with violated distances indicated in red. This highlights the ability of FAIMS-GPF to retrieve XLs of high structural relevance and specificity.

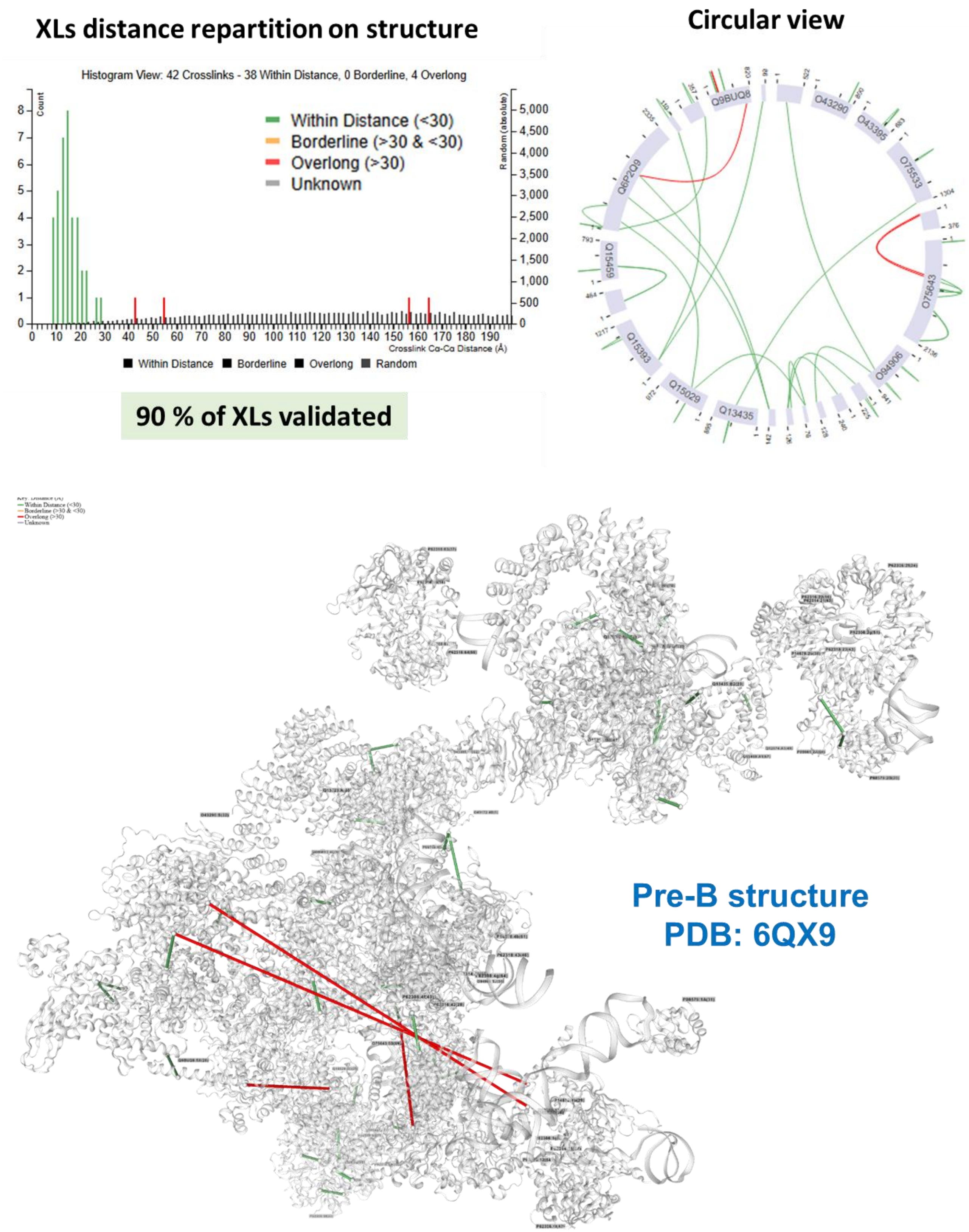


**Figure S19. Distance-validation of reproducible XLs on the pre-B spliceosome structure co-existing in the sample.** Among the 42 XLs mapped on the human pre-Bact-1 structure, 90 % could be distance-validated.

**Supplementary Tables.**

| Site 1 | Protein 1 | Site 2 | Protein 2 |
|---|---|---|---|
| 7 | Cdk7 | 52 | Cdk7 |
| 6 | Cdk7 | 14 | Cdk7 |
| 14 | Cdk7 | 32 | Cdk7 |
| 6 | Cdk7 | 32 | Cdk7 |
| 32 | Cdk7 | 52 | Cdk7 |
| 32 | Cdk7 | 328 | Cdk7 |
| 139 | Cdk7 | 160 | Cdk7 |
| 139 | Cdk7 | 328 | Cdk7 |
| 52 | Cdk7 | 164 | Cdk7 |
| 6 | Cdk7 | 328 | Cdk7 |
| 52 | Cdk7 | 328 | Cdk7 |
| 328 | Cdk7 | 342 | Cdk7 |
| 14 | Cdk7 | 328 | Cdk7 |
| 6 | Cdk7 | 342 | Cdk7 |
| 52 | Cdk7 | 274 | CyclinH |
| 164 | Cdk7 | 78 | CyclinH |
| 328 | Cdk7 | 8 | CyclinH |
| 32 | Cdk7 | 189 | CyclinH |
| 342 | Cdk7 | 8 | CyclinH |
| 6 | Cdk7 | 8 | CyclinH |
| 52 | Cdk7 | 8 | CyclinH |
| 6 | Cdk7 | 33 | CyclinH |
| 6 | Cdk7 | 39 | CyclinH |
| 6 | Cdk7 | 72 | CyclinH |
| 52 | Cdk7 | 72 | CyclinH |
| 52 | Cdk7 | 121 | CyclinH |
| 6 | Cdk7 | 189 | CyclinH |
| 52 | Cdk7 | 189 | CyclinH |
| 6 | Cdk7 | 292 | CyclinH |
| 6 | Cdk7 | 295 | CyclinH |
| 32 | Cdk7 | 295 | CyclinH |
| 52 | Cdk7 | 295 | CyclinH |
| 161 | Cdk7 | 78 | CyclinH |
| 164 | Cdk7 | 189 | CyclinH |
| 32 | Cdk7 | 72 | CyclinH |
| 8 | CyclinH | 239 | CyclinH |
| 274 | CyclinH | 295 | CyclinH |
| 8 | CyclinH | 189 | CyclinH |
| 39 | CyclinH | 295 | CyclinH |
| 8 | CyclinH | 72 | CyclinH |
| 72 | CyclinH | 189 | CyclinH |
| 189 | CyclinH | 274 | CyclinH |
| 8 | CyclinH | 14 | CyclinH |
| 32 | Cdk7 | 70 | Mat1 |
| 8 | CyclinH | 70 | Mat1 |
| 72 | CyclinH | 70 | Mat1 |
| 189 | CyclinH | 70 | Mat1 |
| 274 | CyclinH | 70 | Mat1 |

**Table S1. XLs identified (FT 2/3) after XL-MS experiments on CAK without FAIMS**.

| Site 1 | Protein 1 | Site 2 | Protein 2 |
|---|---|---|---|
| 6 | Cdk7 | 32 | Cdk7 |
| 6 | Cdk7 | 41 | Cdk7 |
| 6 | Cdk7 | 52 | Cdk7 |
| 6 | Cdk7 | 291 | Cdk7 |
| 6 | Cdk7 | 342 | Cdk7 |
| 7 | Cdk7 | 52 | Cdk7 |
| 14 | Cdk7 | 32 | Cdk7 |
| 14 | Cdk7 | 328 | Cdk7 |
| 28 | Cdk7 | 32 | Cdk7 |
| 32 | Cdk7 | 52 | Cdk7 |
| 32 | Cdk7 | 328 | Cdk7 |
| 32 | Cdk7 | 342 | Cdk7 |
| 41 | Cdk7 | 52 | Cdk7 |
| 52 | Cdk7 | 160 | Cdk7 |
| 52 | Cdk7 | 164 | Cdk7 |
| 52 | Cdk7 | 170 | Cdk7 |
| 52 | Cdk7 | 291 | Cdk7 |
| 52 | Cdk7 | 328 | Cdk7 |
| 52 | Cdk7 | 342 | Cdk7 |
| 139 | Cdk7 | 328 | Cdk7 |
| 160 | Cdk7 | 328 | Cdk7 |
| 160 | Cdk7 | 342 | Cdk7 |
| 328 | Cdk7 | 342 | Cdk7 |
| 6 | Cdk7 | 8 | CyclinH |
| 6 | Cdk7 | 14 | CyclinH |
| 6 | Cdk7 | 39 | CyclinH |
| 6 | Cdk7 | 72 | CyclinH |
| 6 | Cdk7 | 189 | CyclinH |
| 6 | Cdk7 | 274 | CyclinH |
| 6 | Cdk7 | 292 | CyclinH |
| 6 | Cdk7 | 295 | CyclinH |
| 32 | Cdk7 | 8 | CyclinH |
| 32 | Cdk7 | 72 | CyclinH |
| 32 | Cdk7 | 189 | CyclinH |
| 32 | Cdk7 | 274 | CyclinH |
| 32 | Cdk7 | 295 | CyclinH |
| 52 | Cdk7 | 39 | CyclinH |
| 52 | Cdk7 | 72 | CyclinH |
| 52 | Cdk7 | 121 | CyclinH |
| 52 | Cdk7 | 189 | CyclinH |
| 52 | Cdk7 | 274 | CyclinH |
| 52 | Cdk7 | 295 | CyclinH |
| 160 | Cdk7 | 72 | CyclinH |
| 164 | Cdk7 | 189 | CyclinH |
| 291 | Cdk7 | 189 | CyclinH |
| 328 | Cdk7 | 8 | CyclinH |
| 342 | Cdk7 | 8 | CyclinH |

| Site 1 | Protein 1 | Site 2 | Protein 2 |
|---|---|---|---|
| 6 | Cdk7 | 70 | Mat1 |
| 6 | Cdk7 | 80 | Mat1 |
| 6 | Cdk7 | 135 | Mat1 |
| 32 | Cdk7 | 70 | Mat1 |
| 52 | Cdk7 | 70 | Mat1 |
| 52 | Cdk7 | 127 | Mat1 |
| 160 | Cdk7 | 70 | Mat1 |
| 291 | Cdk7 | 70 | Mat1 |
| 328 | Cdk7 | 70 | Mat1 |
| 342 | Cdk7 | 70 | Mat1 |
| 8 | CyclinH | 12 | CyclinH |
| 8 | CyclinH | 14 | CyclinH |
| 8 | CyclinH | 189 | CyclinH |
| 8 | CyclinH | 239 | CyclinH |
| 8 | CyclinH | 253 | CyclinH |
| 33 | CyclinH | 295 | CyclinH |
| 39 | CyclinH | 189 | CyclinH |
| 39 | CyclinH | 292 | CyclinH |
| 39 | CyclinH | 295 | CyclinH |
| 72 | CyclinH | 189 | CyclinH |
| 72 | CyclinH | 295 | CyclinH |
| 189 | CyclinH | 274 | CyclinH |
| 189 | CyclinH | 295 | CyclinH |
| 190 | CyclinH | 295 | CyclinH |
| 274 | CyclinH | 295 | CyclinH |
| 8 | CyclinH | 70 | Mat1 |
| 8 | CyclinH | 167 | Mat1 |
| 72 | CyclinH | 70 | Mat1 |
| 189 | CyclinH | 70 | Mat1 |
| 189 | CyclinH | 127 | Mat1 |
| 189 | CyclinH | 167 | Mat1 |
| 274 | CyclinH | 70 | Mat1 |
| 70 | Mat1 | 80 | Mat1 |
| 70 | Mat1 | 176 | Mat1 |
| 127 | Mat1 | 135 | Mat1 |

**Table S2. XLs identified (FT 2/3) after XL-MS experiments on CAK with 4-CV GPF FAIMS method**.
